# RENANO: a REference-based compressor for NANOpore FASTQ files

**DOI:** 10.1101/2021.03.26.437155

**Authors:** Guillermo Dufort y Álvarez, Gadiel Seroussi, Pablo Smircich, José Sotelo-Silveira, Idoia Ochoa, Álvaro Martín

## Abstract

Nanopore sequencing technologies are rapidly gaining popularity, in part, due to the massive amounts of genomic data they produce in short periods of time (up to 8.5 TB of data in less than 72 hours). In order to reduce the costs of transmission and storage, efficient compression methods for this type of data are needed. Unlike short-read technologies, nanopore sequencing generates long noisy reads of variable length. In this note we introduce RENANO, a reference-based lossless FASTQ data compressor, specifically tailored to compress FASTQ files generated with nanopore sequencing technologies. RENANO builds on the recent compressor ENANO, which is the current state of the art. RENANO focuses on improving the compression of the base call sequence portion of the FASTQ file, leaving the other parts of ENANO intact. Two novel reference-based compression algorithms are introduced, contemplating different scenarios: in the first scenario, a reference genome is available without cost to both the compressor and the decompressor; in the second, the reference genome is available *only* on the compressor side, and a compacted version of the reference is transmitted to the decompressor as part of the compressed file. To evaluate the proposed algorithms, we compare RENANO against ENANO on several publicly available nanopore datasets. In the first scenario considered, RENANO improves the base call sequences compression of ENANO by 39.8%, on average, over all the datasets. As for total compression (including the other parts of the FASTQ file), the average improvement is 12.7%. In the second scenario considered, the base call compression improvements of RENANO over ENANO range from 15.2% to 49.0%, depending on the coverage of the compressed dataset, while in terms of total size, the improvements range from 5.1% to 16.5%.

Implementations of the proposed algorithms are freely available for download at https://github.com/guilledufort/RENANO.

## 1 Background

We investigate compression of DNA data generated by *nanopore sequencing technologies* [29] *and stored in the FASTQ format [3]. A FASTQ file stores the result of a sequencing process, which consists of a set of readings of genome fragments, called reads*. Each read contains an *identifier string*, a *base call sequence* (also referred to as read), and a *quality score sequence*. The identifier string is generally a short free text segment, which identifies the read. The base call sequence is a string of letters from the set {*A, C, G, T, N*}, where letters *A, C, G*, and *T*, represent the nitrogenous bases (base-pairs) of a DNA sequence, and letter *N* is a special character that represents a nitrogenous base that could not be determined. For convenience, we refer to the base call sequence as a *base call string*, as it will often be used as a string throughout the rest of this document. Finally, the quality score sequence is a string of symbols, of the same length as the base call string, where the *i*-th symbol encodes an estimated probability of the *i*-th base call being correct.

The characteristics of the base call string and the quality score sequence that compose each read vary depending on the technology used to perform the DNA sequencing. For example, Second Generation Sequencing (SGS) technologies generate high quality short reads (a few hundred base-pair long), generally of fixed length, with a quality score alphabet that ranges from 4 values to about 40, depending on the specific technology. On the other hand, nanopore technologies generate very long reads (up to hundreds of thousands base-pair long [25]), of variable length, which generally have lower quality than those generated by SGS, and a larger quality scores alphabet size (94 for nanopore).

In a previous work we developed ENANO [5], a lossless compression algorithm especially designed for nanopore FASTQ files. ENANO achieves state of the art compression performance by using a novel quality score sequence compression algorithm that exploits the specific characteristics of nanopore data. In this work we focus on further improving the compression of nanopore FASTQ files by addressing, specifically, the compression of the base call strings.

Many algorithms have been proposed in the literature to compress the base call strings of a FASTQ file [7]. These compression algorithms can be roughly divided into two categories: *reference-free* methods, in which no external information is used to aid compression, and *reference-based* methods, in which an external reference genome is used to aid compression. A reference genome file stores the base call strings that compose the chromosomes of a genome, usually in FASTA format [21], which is a variation of the FASTQ format that does not store quality scores.

Reference-free compression methods have the advantage of being self-contained and not needing any external resource to work properly, but usually lack in compression performance compared to reference-based methods. On the other hand, reference-based methods require access to a proper reference genome file, that is, a genome of the same or similar species as the one sequenced and stored in the FASTQ file. In turn, these methods achieve better compression performance (see, e.g., [12], [8], [2]) by exploiting the similarities between the sequenced and the reference genomes, which, for example, in the case of human genomes exceeds 99% of the base-pairs [17].

To exploit the information provided by an external reference genome, reference-based compression algorithms start by *aligning* the base call strings of the FASTQ file against the strings in the reference genome file, producing a series of *alignments*. Loosely speaking, an alignment is a description of a string *q* in terms of a reference string *r*, which describes the editing operations needed on the reference string *r* to produce the string *q* (these notions, as applied in this work, will be defined more precisely in Section 2.1). If the aligned string *q* is similar to the reference string *r*, then the alignment serves as a compact representation of *q*. Consequently, reference-based compression algorithms efficiently compress *q* by encoding the alignment instead. The decoder can later reconstruct the string *q* by decoding the alignment and applying the editing operations described in it to the reference string *r*, which is assumed available on the decoder side.

The availability of a proper reference genome is not an uncommon scenario in bioinformatics. In fact, many bioinformatic analysis tasks performed on FASTQ data, such as sequence analysis or gene expression analysis, already require a step in their pipeline where the reads of the FASTQ file are aligned against a reference genome [19], using specialized alignment tools. Even for metagenomic or contaminated samples, where the sequenced organisms may not be known in advance, an appropriate reference can be readily obtained by concatenating the genomes of the most prevalent species identified by a taxonomic classification tool [26, 30, 13] (see Section 3.1). In the case of the long noisy reads produced by nanopore technologies, the state of the art alignment tool is Minimap2 [20], which outputs a file in *Pairwise mApping Format* (PAF) with the result of aligning each read of a FASTQ file against a reference genome. In this work, we focus on exploiting these scenarios by taking a reference-based approach to compressing the read base call strings of the FASTQ file.

As mentioned, the decoding process requires access to the same reference genome file used during compression, which may not always be practical. To address this problem, reference-based compression methods can encode the relevant parts of the reference genome file into the compressed file, thus sacrificing some compression performance in exchange for making the decompression algorithm independent of the original reference genome.

Most of the reference-based compression algorithms for base call strings available in the literature [15, 8, 2, 12, 14, 1, 6, 9] are specifically tailored to compress high quality short reads generated by SGS technologies, and therefore do not perform well (or at all) on the noisy long reads generated by nanopore technologies. The more recent work [16] presents a compression tool capable of compressing nanopore FASTQ files, which offers both a reference-free mode and a reference-based mode. However, our experiments have shown that, when running both methods on nanopore FASTQ data, the reference-free mode consistently outperforms the reference-based mode, and the reference-based mode fails to compress some of the tested datasets (results of these experiments are presented in Section 3.4.2).

In this note we introduce RENANO, a lossless nanopore FASTQ data compressor that builds on its predecessor ENANO, introducing two novel reference-based compression algorithms for base call strings that significantly improve the state of the art compression performance. The working principle of both algorithms is the same, with the difference being that one assumes the reference genome to be available at the decoder, whereas the other one encodes the parts of the reference genome needed at the decoder as part of the compressed output.

The ENANO compressor [5] parses the input FASTQ file into blocks, whose boundaries are dynamically determined so that every read is entirely contained in a single block. Each block is divided into three streams: read identifiers, base call strings, and quality score sequences. For each stream ENANO uses a specific *context model* [27] to determine a probability distribution for each symbol, which, in turn, is passed to an arithmetic encoder [28] to generate the output bit stream. These probability distributions are dynamically updated as the file is sequentially compressed. The decompressor works in lockstep, symmetrically. The input file is encoded while the statistics are adaptively updated for each context model, for a certain number of blocks of the file. Afterwards, the statistics are fixed, allowing for fast parallel compression and decompression. Full details are given in [5]. We build RENANO by replacing the base call strings compression scheme of ENANO with one of two novel reference-based compression algorithms introduced in Sections 2, denoted by RENANO_1_ and RENANO_2_, while keeping the processing of the other streams intact. The two variants of RENANO thus constructed are complete FASTQ file compression schemes. However, since the rest of the paper deals only with the compression of base call strings, we will slightly abuse terminology and use RENANO_1_ and RENANO_2_ also to refer specifically to the compression algorithms for base call strings. For RENANO_1_, which we present in Section 2.2, we assume that a reference genome is available without cost to both the compressor and the decompressor. On the other hand, RENANO_2_, presented in Section 2.3, does not require a reference genome on the decompression side.

The rest of the note is organized as follows. Section 2 introduces basic notation, terminology, and tools and, as mentioned, describes the two proposed reference-based base call string compression algorithms. Section 3 presents experimental results, and comparisons of RENANO to ENANO and to the scheme Genozip of [16]. The experiments reveal that both variants of RENANO significantly outperform the schemes they are compared against.

## 2 Methods

In this section we present RENANO_1_ and RENANO_2_, two novel long-read base call string compression algorithms that use the alignment information (against a reference sequence) to improve compression. The general idea behind the proposed compression schemes is to encode a large portion of each base call string as a series of alignments to a reference genome, such that the alignments can be described more compactly than the raw base call string, thus yielding better compression. For both schemes, we assume that a reference sequence (e.g., a genome) in FASTA format is available on the compressor side. For RENANO_1_, we consider the scenario in which the reference sequence is also available without cost to the decompressor, while for RENANO_2_ we consider the scenario in which the reference sequence is *not* available to the decompressor, but a compacted version of that sequence is stored as part of the compressed output, thus incurring a code length cost. In both cases, we assume that the alignment information is obtained from an available alignment tool, such as Minimap2 [20], which generates a file in PAF format. Note that even though the current implementation of our compressor expects a PAF file as input, the algorithm could readily be adapted to other formats if they become popular in the future. Finally, both algorithms assume the base call strings are stored in the widely used FASTQ format [3]. We show in Section 3 that both compression schemes exhibit improvements in compression ratio as compared to ENANO [5], which does not use a reference sequence or alignment information, and to Genozip [16], which offers two compression modes: a reference-based mode, and a reference-free mode.

Both of the presented compression algorithms take the alignment information generated by the alignment tool and transform it to an internal representation, which, together with the reference sequence, allows for perfect reconstruction of the original base call strings. The ability to encode the internal representation of alignments compactly is at the core of the schemes. In Section 2.1 we present the general notations and definitions needed to formalize the proposed internal alignment representation. Sections 2.2 and 2.3 describe RENANO_1_ and RENANO_2_, respectively, in detail. Finally, Section 2.4 provides specific details on how the PAF formatted output Minimap2 [20] is transformed into our internal representation.

Implementations of full FASTQ file compressors/decompressors based on RENANO_1_ and RENANO_2_ are freely available for download at https://github.com/guilledufort/RENANO.

### 2.1 Notations and definitions

Let Σ = {*A, C, G, T, N}* be the alphabet of base calls, where {*A, C, G, T}* represent the DNA nucleotides, and *N* represents an undetermined base call. For a symbol *b* ∈ Σ, 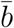 is its complementary nucleotide, that is, *Ā* = *T*, 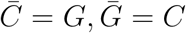, and 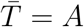, and for symbol N we define 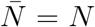. We say that a string *s* = *s*_1_*s*_2_…*s*_*n*_, *s*_*i*_ ∈ Σ, is a *base call string* of length |*s*| = *n*, and define its *reverse complement* as 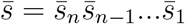. We also use the notation *s*[*i* : *j*] = *s*_*i*_…*s*_*j*_, to denote a substring of length *j* − *i* + 1, starting at position *i* and ending at position *j*; if *j < i*, we let *s*[*i* : *j*] be the empty string, denoted by *λ*. For a base call string *s* and *d* ∈ {0, 1}, we define the *strand function π*(*s, d*) as *π*(*s, d*) = *s* if *d* = 0, and 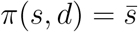 if *d* = 1. We refer to *d* as the *strand direction indicator* of *π*(*s, d*).

We define an *encoding transformation, ϕ*, as a sequence of *K* + 1 *transformation steps* that converts a base call string, *s*, into another base call string, *ϕ*(*s*), where *K* is a positive integer. The first *K* transformation steps construct a string *s′*, starting from *s′* = *λ*, while scanning two input strings: *s*, and an additional (given) string ℐ. Each such transformation step is represented by a triplet of non-negative integers, (*I, S, M*), that can be interpreted as driving three elementary string operations on *s′*, in order:

1. append a copy of the next *I* symbols from string ℐ to *s′*;
2. skip the following *S* symbols of *s*;
3. append a copy of the next *M* symbols from *s* to *s′*.

The (*K* + 1)-th, and final, step of the encoding transformation consists of applying the strand function to the constructed string *s′*, obtaining the result *ϕ*(*s*) = *π*(*s′, d*), given a strand direction indicator value *d*. More specifically, we let *ϕ* = ({(*I*_*k*_, *S*_*k*_, *M*_*k*_)}_1≤*k*≤*K*_, ℐ, *d*), where each triplet (*I*_*k*_, *S*_*k*_, *M*_*k*_) is comprised of an *insertion* length *I*_*k*_, a *skip* length *S*_*k*_, and a *match* length *M*_*k*_; ℐ is the *insertions base call string* of *ϕ*, with length 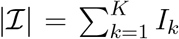; and *d* is the strand direction indicator.

We always apply *ϕ*(*s*) to strings *s* of length equal to 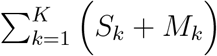. Algorithm 1 summarizes the encoding transformation process. An example of the application of Algorithm 1 is shown in Figure 1.

**Figure 1:**
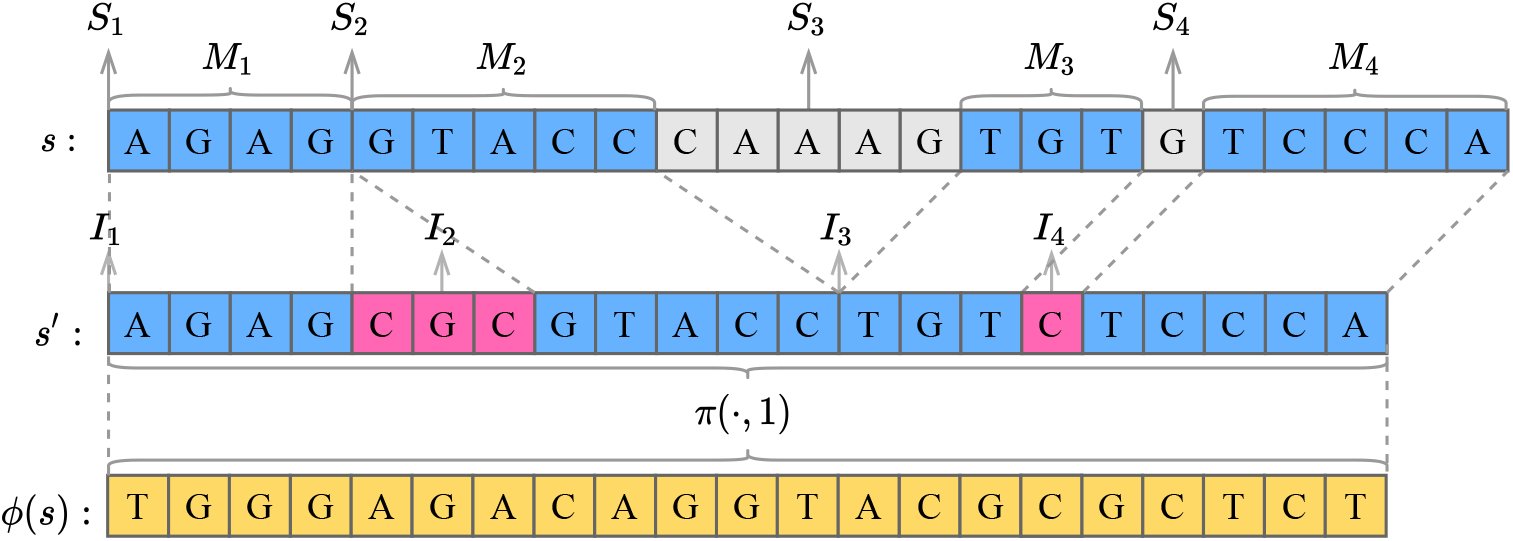
Constructing *ϕ*(*s*) by executing Algorithm 1 with inputs: *s*, and encoding transformation *ϕ* = (((0, 0, 4), (3, 0, 5), (0, 5, 3), (1, 1, 5)), ℐ = CGCC, *d* = 1). Notice that when an insertion length or a match length is 0, no symbols are appended to *s′* from ℐ or *s*, respectively. Also, if the strand direction indicator is *d* = 0, then *ϕ*(*s*) = *s′*.

A reference genome is usually composed of multiple base call strings; for example, in the case of the human genome there is at least one base call string for each chromosome. Therefore, we represent a reference genome as an ordered set, ℛ = *r*_*k* 1≤*k*≤|ℛ|_, of base call strings. Now, let *q* be a read base call string, and let *r* ∈ ℛ be a reference base call string. We define an *atomic alignment* between *q* and *r*, denoted *α*(*q, r*), as an encoding transformation from a substring of *r* to a substring of *q*. More specifically, we let *α*(*q, r*) = (*i, j, i′, j′, ϕ*), where *ϕ* is an encoding transformation such that *ϕ*(*r*[*i′* : *j′*]) = *q*[*i* : *j*]. Finally, we define a *full alignment* between *q* and ℛ, as a sequence of atomic alignments 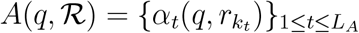, where *k*_*t*_ is the index of a reference string in ℛ, and *L*_*A*_ is the length of (i.e., number of atomic alignments in) *A*(*q*, ℛ). We consistently use the same subscript *t* of an atomic alignment 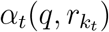 to denote all of its components, i.e., 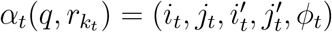. In Figure 2 we show an example of a full alignment. We refer to a full alignment *A*(*q, ℛ*) as *non-overlapping*, if none of the aligned substrings of *q* overlap (such as the one shown in Figure 2). Notice that the atomic alignments in the example do not cover the whole read *q*, a situation that may also occur in the general case. We refer to segments of *q* not covered by *A*(*q, ℛ*) as *unaligned*. Later on, we will discuss how these segments are dealt with in our algorithms.

**Figure 2:**
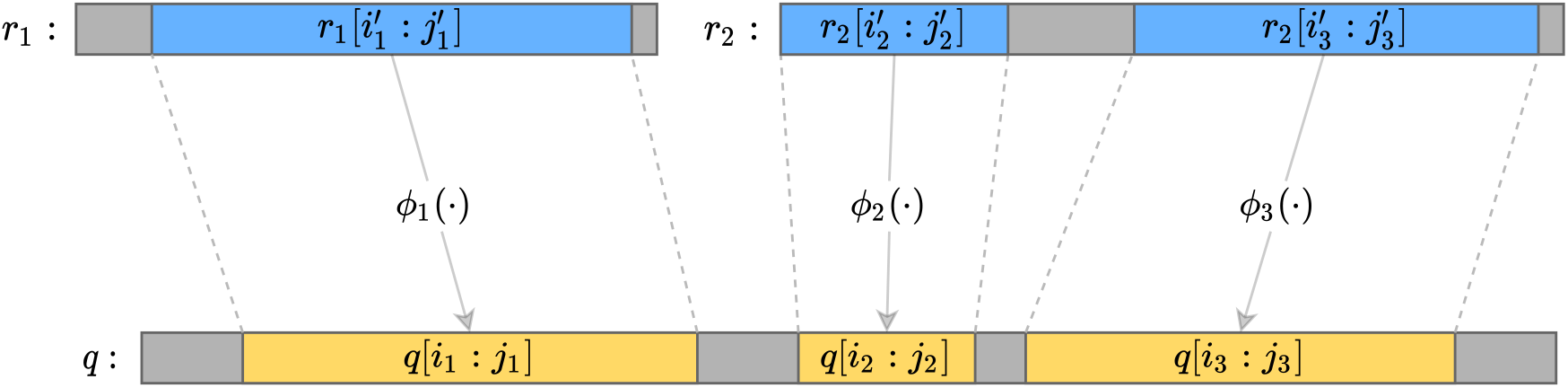
Example of a full alignment *A*(*q*, ℛ) = {*α*_1_(*q, r*_1_), *α*_2_(*q, r*_2_), *α*_3_(*q, r*_2_)}, between a base call string *q* and a set of reference base call strings ℛ = {*r*_1_, *r*_2_}.

#### Algorithm 1: Convert string *s* into string *ϕ*(*s*).

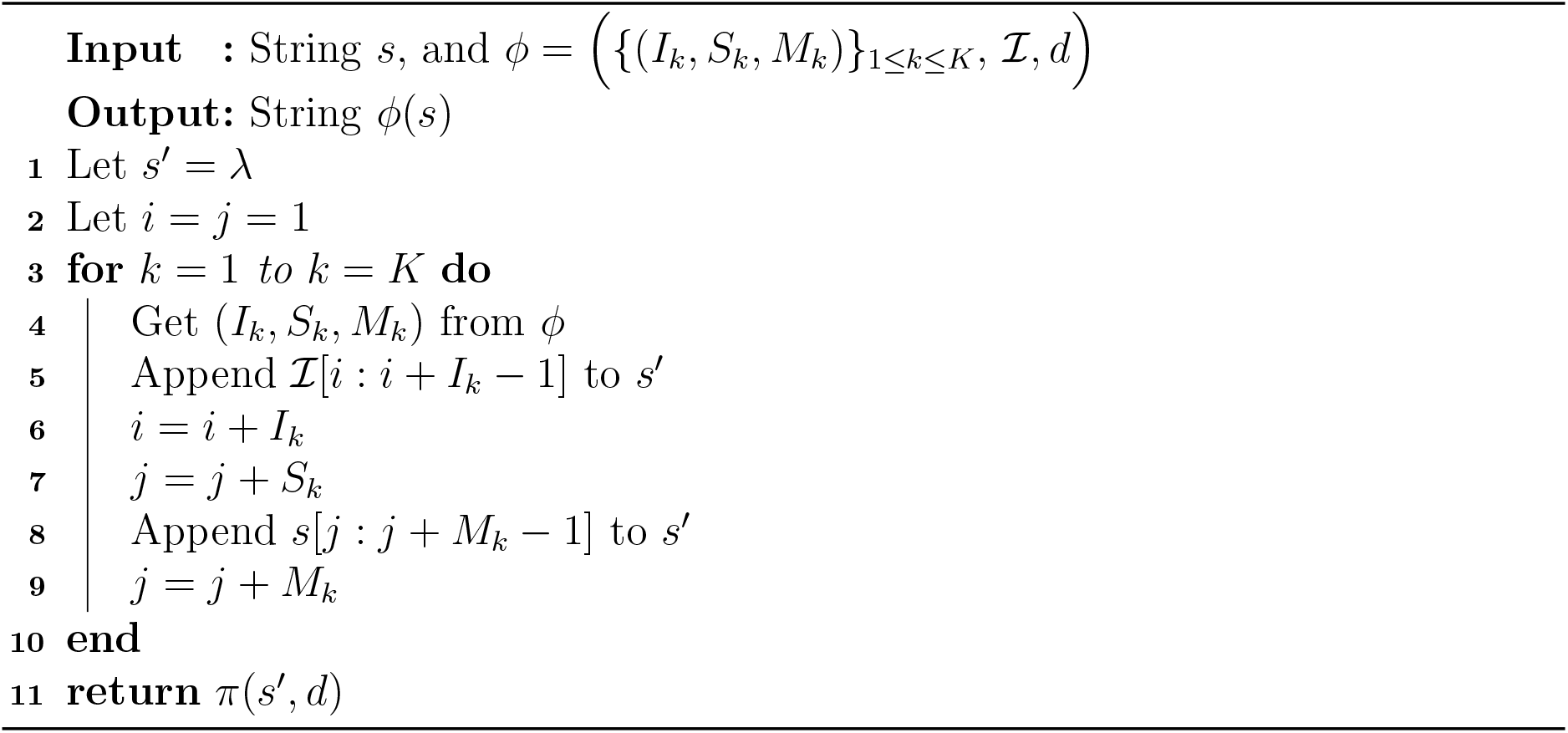

### 2.2 RENANO_1_: A reference-dependent compression and decompression scheme

In this section we present RENANO_1_, an algorithm that compresses the base call strings of a FASTQ file assuming that the set of reference strings ℛ is available to both the compressor and the decompressor. We further assume that the full alignment *A*(*q, ℛ*) for each base call string *q* against the set of reference strings ℛ is available during the compression stage. In practice, this means that we assume an alignment tool has been run on the base call strings in the file against the set of reference strings ℛ, and that *A*(*q, ℛ*) has been derived from the alignment information obtained (details on how this is done are provided in Section 2.4). For convenience, we also assume the atomic alignments in 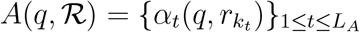 to be non-overlapping (as exemplified in Figure 2). Again in Section 2.4, we show that any full alignment can be transformed to satisfy this assumption.

As discussed in Section 1, RENANO builds over its predecessor ENANO [5], which parses and compresses the FASTQ file in blocks that always contain a whole number of reads. For consistency, our algorithm compresses the sequence of read base call strings contained in a block of the FASTQ file, denoted by *Q* = *q*_1_, *⋯*, *q*_|*Q*|_, independently of the base call strings contained in other blocks. This makes the compression scheme compatible with the parallelization strategy implemented in ENANO.

Recall from Section 2.1 that, given an atomic alignment *α*_*t*_(*q, r*) = (*i, j, i ′, j ′, ϕ*), we can reconstruct the substring *q*[*i* : *j*] = *ϕ*(*r*[*i ′, j ′*]) by executing Algorithm 1 with inputs *ϕ* and *r*[*i ′, j ′*]. As we have access to the reference string *r* during decompression, encoding the atomic alignment *α*_*t*_(*q, r*) suffices to describe *q*[*i* : *j*]. Extending this idea, RENANO_1_ compresses the aligned substrings of a read base call string *q* by encoding the full alignment *A*(*q, ℛ*), while the unaligned parts of *q* are encoded as raw base call strings, as in ENANO. We now proceed to describe the compression scheme in detail.

#### 2.2.1 The encoding algorithm

To encode *Q*, we split the various elements of the representations of its read base call strings *q*_*i*_, i.e, the aligned substrings represented in *A*(*q*_*i*_, *ℛ*) and the unaligned substrings, into separate sequences, which we call *streams*. We refer to the stream values that comprise a representation of any of these elements as its *stream representation*. To help improve compression performance, the streams are designed so that each stream gathers parts of the representations that we expect to be correlated. The streams, in turn, are encoded independently. Different streams may contain different types of data. In particular, as described in more detail below, we will have streams comprised of raw base call symbols, streams comprised of binary symbols, and streams comprised of non-negative integers. For each stream *S* of the latter, we define a parameter *η*_*S*_ that determines the bit size representation of the integers, where *η*_*S*_ is, for convenience, assumed to be a positive multiple of 8. The values of these parameters used in our experiments are discussed later in Section 3.3. Specifically, we define the following streams:

- ℬ: base call strings stream used to encode individual raw base call symbols (A, C, G, T, N), which include: the unaligned base call strings, and the insertion base call strings ℐ that are part of the encoding transformations.
- ℒ: base call string lengths stream used to store the non-negative integer |*q*|, i.e., the length of each read basecall string *q*, using an *η* _ℒ_-bit representation.
- 𝒬: full alignment sizes stream used to store the non-negative integers *L*_*A*_, using an *η* _𝒬_-bit representation.
- 𝒮: starting position increments stream for aligned substrings of *q*, i.e., the non-negative integers *i*_*t*_ − *i*_*t*−1_, with the convention *i*_0_ = 0, using an *η* _𝒮_ -bit representation.
- ℰ: lengths stream for aligned substrings of *q*, i.e., the non-negative integers *z*_*t*_ = *j*_*t*_ − *i*_*t*_ + 1, using an *η* _ℰ_ -bit representation.
- 𝒰: aligned reference string identity indexes stream used to store the non-negative integers 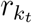, using an *η* _𝒰_ -bit representation.
- 𝒮 ′: aligned reference substring starting positions stream, which stores the non-negative integers 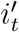, using an *η* _𝒮 ′_-bit representation.
- 𝒟: strand direction indicator stream, which stores the binary values of *d*_*t*_ ∈ {0, 1}.
- 𝒩: insertion lengths stream that stores the non-negative integers *I*_*k*_, using an *η* _𝒩_ -bit representation.
- 𝒦: skip lengths stream, which stores the non-negative integers *S*_*k*_, using an *η* _𝒦_-bit representation.
- ℳ: match lengths stream used to store the non-negative integers *M*_*k*_, using an *η* _ℳ_-bit representation.

For convenience, we define 𝕊 = {ℬ, *ℒ*, *𝒬, 𝒮, ℰ*, *𝒰*, *𝒮′, 𝒟, 𝒩, 𝒦*, *ℳ*} as the set of all the proposed streams.

##### Algorithm 2: Encode a sequence of read base call strings.

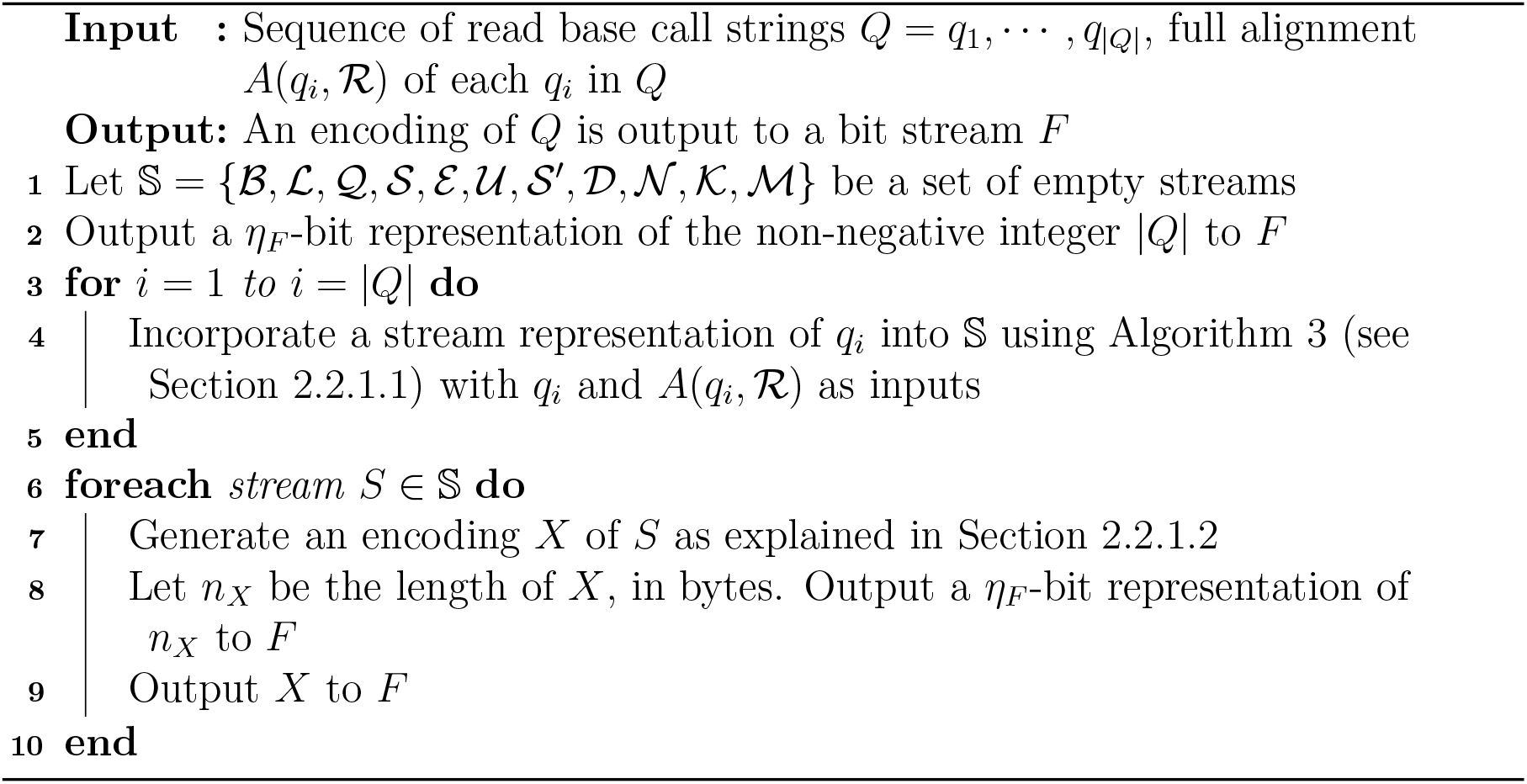

In Algorithm 2 we present the general compression scheme for a sequence of base call strings *Q* = *q*_1_, …, *q*_|*Q*|_. The algorithm receives *Q* as input together with a full alignment *A*(*q*_*i*_, *ℛ*) for each *q*_*i*_ in *Q*, and generates an encoding of *Q* that is output to a bit stream *F*. All the streams in 𝕊 are initialized as empty sequences in Step 1. In Step 2, the algorithm outputs the number of read base call strings in *Q*, so that the decoder can retrieve how many read base call strings are encoded into the streams. In Step 3, the algorithm loops over each base call string *q*_*i*_, splitting a representation of *q*_*i*_ into the various streams in 𝕊 by running Algorithm 3, which we discuss in Subsection 2.2.1.1. Then, the loop in Step 6 encodes the content of each stream *S* in 𝕊. To this end, in Step 7, *S* is encoded using compression techniques that we present later in Subsection 2.2.1.2. Next, the encoding of the stream is output to *F* preceded by its length (in bytes). This ensures that the decoder can access each stream separately. Both numbers output in steps 2 and 8 are encoded as *η*_*F*_ -bit integers, where *η*_*F*_ is an implementation parameter discussed in Section 3.3. We now proceed to describe steps 4 and 7 of the algorithm in detail.

##### 2.2.1.1 Splitting the representation of each base call string into separate streams (Step 4 of Algorithm 2)

To generate a stream representation of an individual read base call string *q*, we use Algorithm 3.

The algorithm starts by appending the length of *q* to stream ℒ (Step 1), and the number of atomic alignments in *A*(*q, ℛ*) to stream 𝒬 (Step 2). A representation of *q* is incorporated into the streams progressively, scanning *q* from start to end as the algorithm iterates over the atomic alignments in *A*(*q*, ℛ), which are previously sorted in increasing order of *i*_*t*_ (Step 3). The variable *i*, initialized to *i* = 1 in Step 4, maintains the starting position of the portion of *q* that remains to be incorporated into the stream representation. In Step 5, the algorithm loops over each atomic alignment in *A*(*q, ℛ*). Each iteration starts by checking, in Step 6, if the current value of *i* is smaller than the starting position of the current atomic alignment, *i*_*t*_. If so, then *q*[*i* : *i*_*t*_ − 1] is an unaligned substring of *q*, which is directly appended to stream ℬ in Step 7. The index *k*_*t*_ of the reference string of the current atomic alignment, 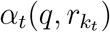, is appended to the stream 𝒰 in Step 10, followed by an execution of Algorithm 4, which appends a representation of 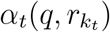 itself to the streams in 𝕊. The iteration ends by updating the value of *i* to the next position yet to be represented, *i* = *j*_*t*_ + 1. By the end of the loop in Step 5, every symbol in *q* has been represented as part of either a raw base call string in Step 7 or an atomic alignment in Step 11, with the possible exception of the trailing symbols in an unaligned suffix of *q*, which are incorporated in Step 15.

Notice that the starting and ending positions of the unaligned substrings are not explicitly described, as they are fully determined by the starting and ending positions of the atomic alignments, together with the total length of *q*.

To generate a stream representation of each atomic alignment 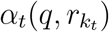, we use Algorithms 4 and 5.

###### Algorithm 3: Describe a full alignment *A*(*q, ℛ*) and the needed parts of base call string *q* using the streams in 𝕊.

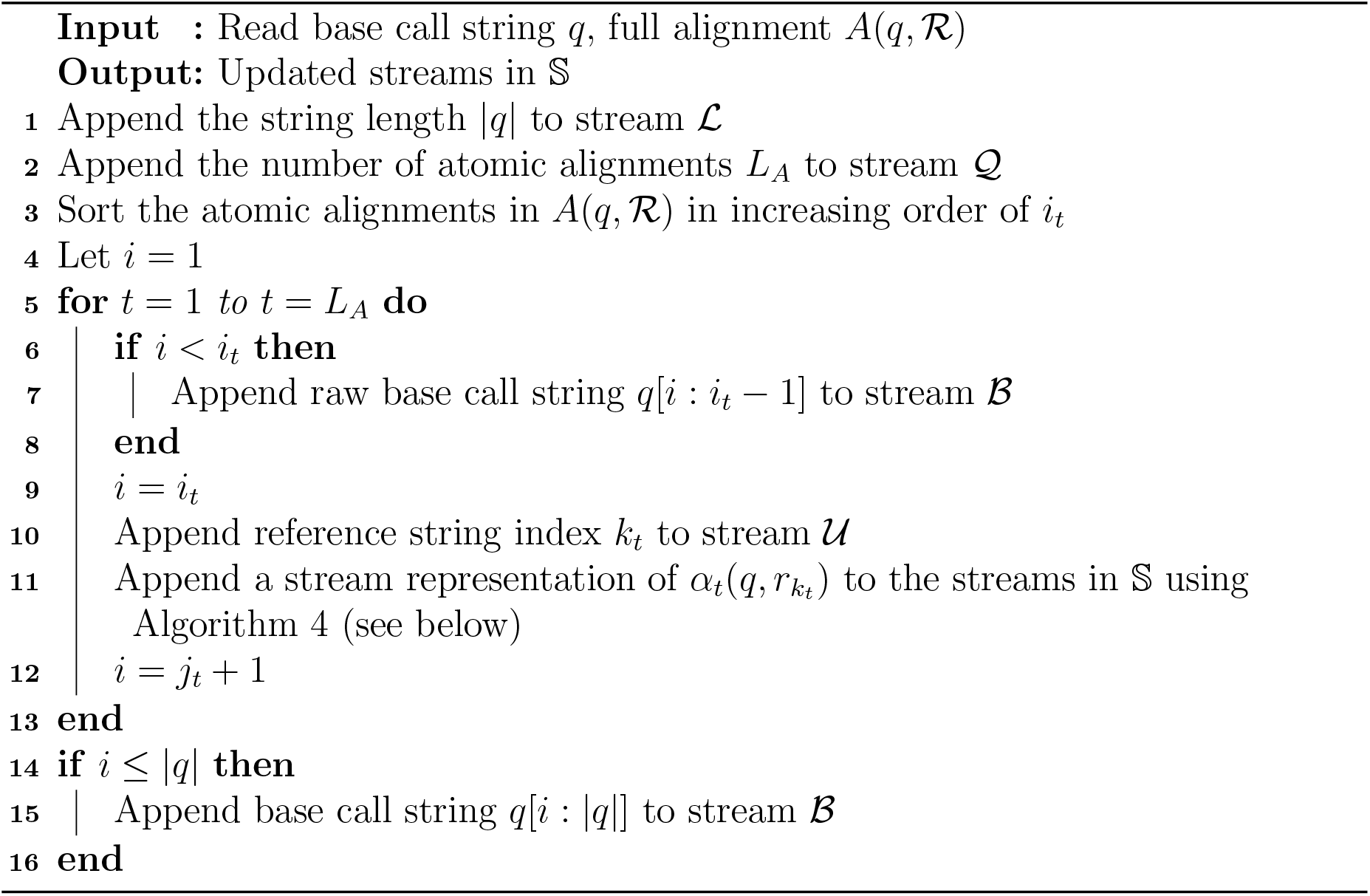

Algorithm 4 starts by appending a differential representation of the starting position of the aligned substring, *i*_*t*_, to stream 𝒮 (Step 2). With this representation, the positions *i*_*t*_ are implicitly determined by the successive differences, *i*_*t*_ − *i*_*t*−1_, which are stored in 𝒮 (recall that by convention, *i*_0_ = 0, and hence the difference is well defined for all *t* ≥ 1). Since *A*(*q, R*) is sorted in increasing order of *i*_*t*_, these differences are non-negative and, in general, with a high frequency of relatively low values, which we exploit to achieve efficient compression. In Step 3, the length of the aligned substring, *z* = *j*_*t*_ − *i*_*t*_ + 1, is appended to stream ℰ. We choose to store the length of the substring instead of its ending position, as this also generally results in a high frequency of relatively low values. Step 4 appends the starting position of the reference substring, 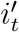, to stream 𝒮′, and the following steps generate a stream representation of the encoding transformation *ϕ*_*t*_. In Steps 5 and 6, the insertions base call string ℐ and the strand direction indicator *d* are appended to streams ℬ and 𝒟, respectively. We do not store the length of the insertions base call string ℐ,as it can be calculated as the sum of the insertion lengths,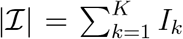. Finally, in Step 7, the algorithm loops through every triplet of the encoding transformation, and incorporates its stream representation into 𝕊 using Algorithm 5.

###### Algorithm 4: Generate a stream representation of an atomic alignment.

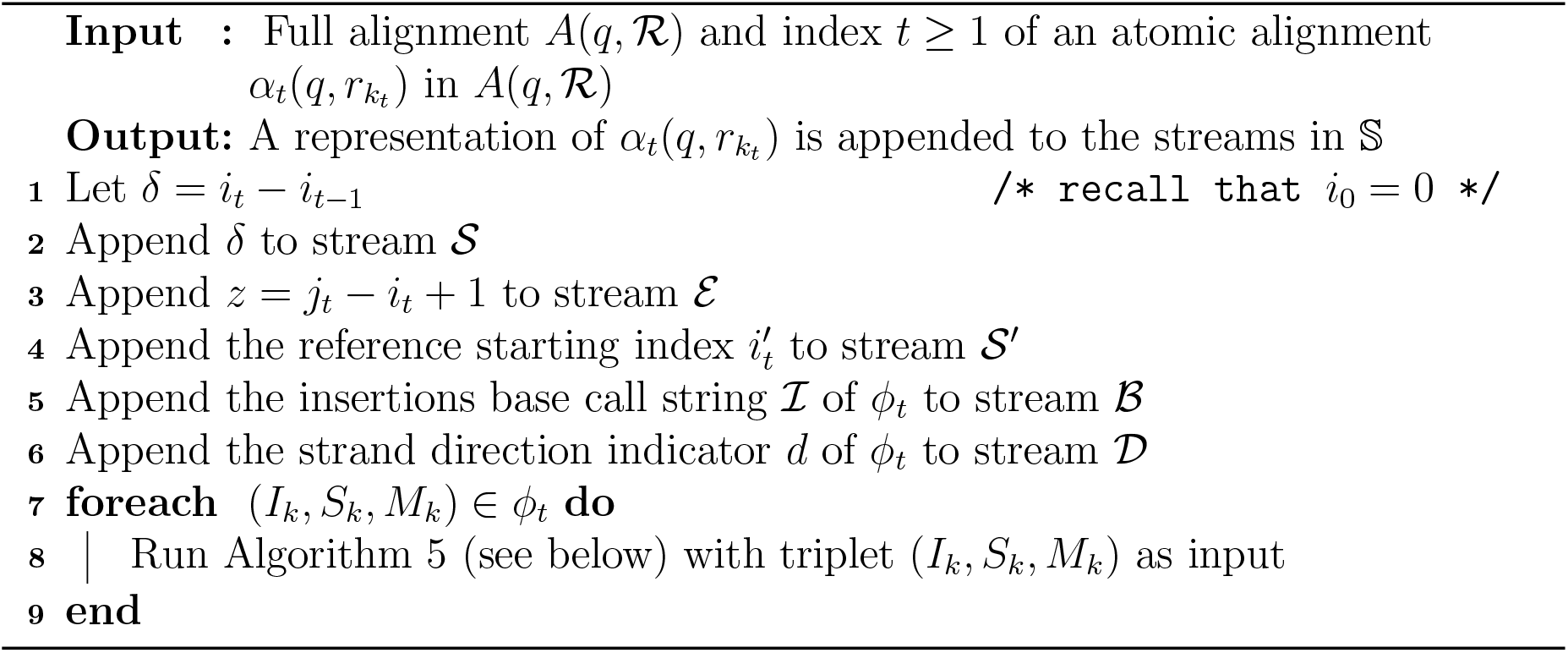

We observed, empirically, that most of the values that typically compose triplets (*I, S, M*) lie within a narrow range of small non-negative integers, with, possibly, a few outliers. For this reason, it is convenient to choose small values for the bit size parameters *η*_𝒩_, *η*_𝒦_, and *η*_ℳ_, used to represent *I, S*, and *M* in the streams 𝒩, 𝒦, and ℳ, respectively. These parameters, in turn, determine thresholds, *T*_𝒩_ = 2^*η 𝒩*^ − 1 for insertion lengths, *T*_𝒦_ = 2^*η 𝒦*^− 1 for skip lengths, and *T*_ℳ_ = 2^*η ℳ*^− 1 for match lengths. Since a stream *S*, for *S* ∈ {𝒩, 𝒦, ℳ}, does not admit a single-value representation of an operation length larger than the threshold *T*_*S*_, every triplet (*I, S, M*) with such an operation length is divided by Algorithm 5 into a sequence of triplets, with trimmed operation lengths that do satisfy these constraints. This substitute sequence represents a combination of string operations that produce the same result as the original operations represented by (*I, S, M*).

Algorithm 5 proceeds recursively. If an operation length, say *I*, is larger than the corresponding threshold, *T*_𝒩_, then the algorithm is recursively invoked for two triplets. One of them represents a single insertion of maximal length, *T*_𝒩_. The other represents the remaining operations, in this case an insertion of length *I* − *T*_𝒩_ together with the original skip and match operations, of lengths *S* and *M*, respectively. Skip and match lengths exceeding the thresholds *T*_𝒦_ and *T*_ℳ_, respectively, are handled analogously.

###### Algorithm 5: Recursively generate a stream representation of a triplet (*I, S, M*), with constrained-length operations.

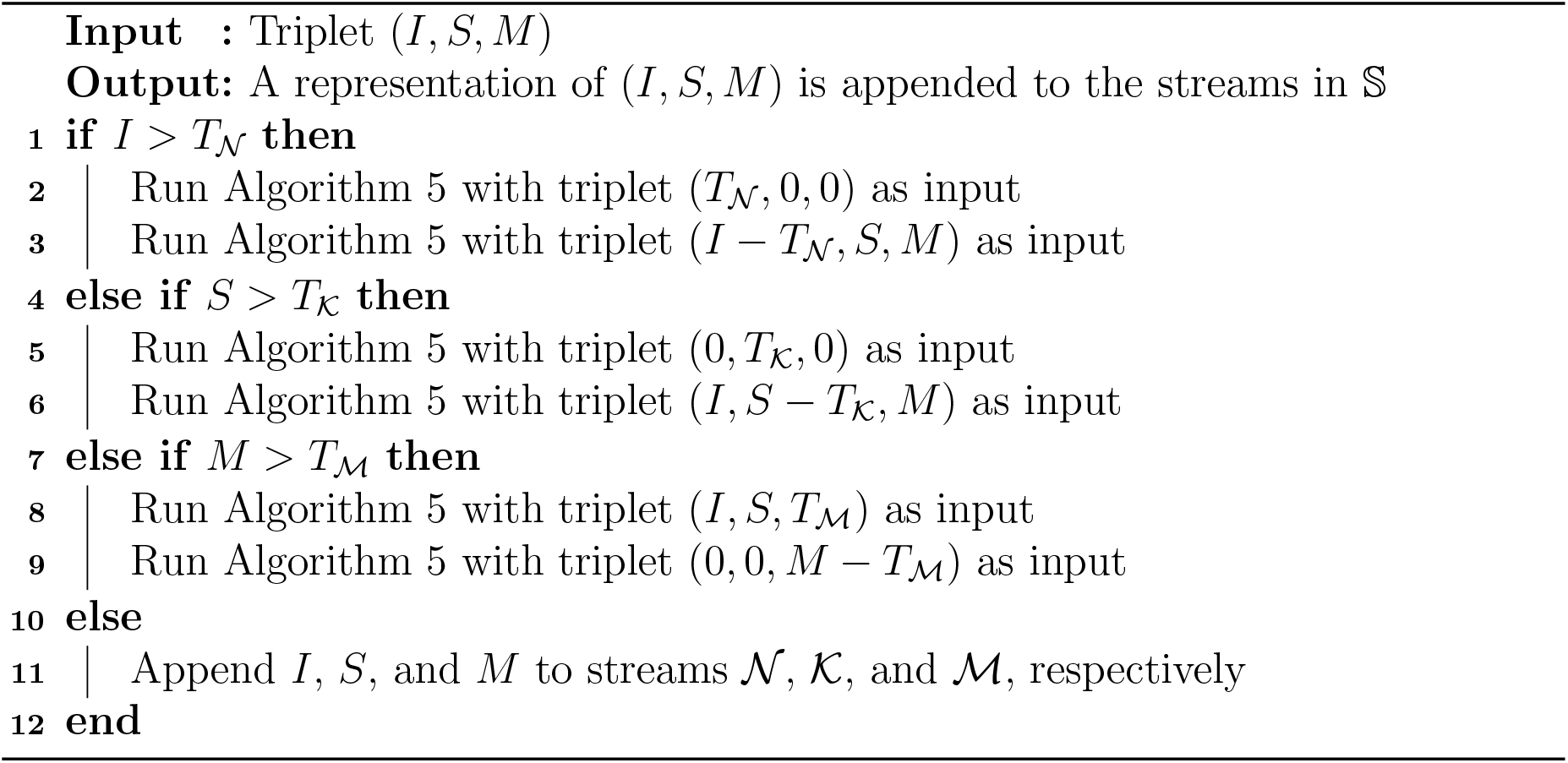

Since an invocation of the algorithm for triplets (*I, S, M*) having at least one value exceeding the corresponding threshold produces recursive calls for triplets (*I′, S′, M ′*), where at least one of *I′, S′, M ′* is strictly smaller than *I, S, M*, respectively (and none of them is larger), then eventually no operation length exceeds its threshold, and the algorithm generates a stream representation of (*I, S, M*) directly in Step 11, with no further recursive calls.

As a last comment regarding Algorithm 4, notice that 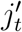 is not explicitly represented in any stream. Nevertheless, the length, *z′*, of the reference substring is implicitly determined by the skip and match lengths of the encoding transformation,

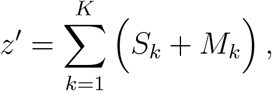

where *K* is the number of transformation steps. The value of 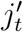, in turn, can be calculated as 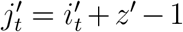. We also notice that the number of transformation steps, *K*, is not explicitly represented either; we show later in Section 2.2.2.1 that it can be reconstructed by the decoder from the streams in 𝕊.

##### 2.2.1.2 Encoding the streams (Step 7 of Algorithm 2)

The algorithms described so far construct a set 𝕊 of streams of various data types, from which, given the reference strings ℛ, the base call strings in the original FASTQ file can be fully reconstructed. To complete a compression algorithm, we must encode these streams, as efficiently as possible, into bitstreams.

To encode each stream we follow a method similar to ENANO [5], which combines *context modeling* [27] with *arithmetic encoding* [28]. In short, we encode the sequence of symbols of a stream by sequentially estimating, for each position of the sequence, a probability distribution over the set of possible values for the symbols in the stream (its *alphabet*). Each symbol position is assigned a specific *context*, which is an abstract class that groups symbols that we model as obeying the same probability distribution. The context assigned to each position is selected from a finite set of possible contexts, based on the values of previously encoded symbols, and the probability distribution determined by each context is sequentially and adaptively estimated from statistics collected from previous occurrences of the same context. To encode each symbol, both the symbol and the estimated probability distribution for the position of the symbol are given as input to an arithmetic encoder. The arithmetic encoder, in turn, generates a sequence of bits that encodes the provided sequence of symbols. The overall code length, in bits, for an input sequence *s*_1_, *s*_2_, …, *s*_*n*_ of stream symbols is

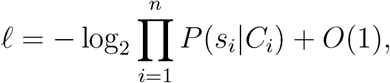

where *C*_*i*_ denotes the context assigned to location *i* in the sequence, and *P* (*s*_*i*_|*C*_*i*_) is the probability assigned to *s*_*i*_ in this context. This code length is, in a well-defined sense, optimal up to an additive constant [28]. On the decoder side, the symbol sequence can be perfectly decoded by calculating the same sequence of probability distributions. Notice that this is possible because the probability distributions are estimated only from occurrences of previously encoded/decoded symbols. We refer the reader to Section 2 of [5] for a more detailed explanation of the method. For the proposed algorithm, we define a specific context model for each stream, as discussed below.

To encode a base call symbol from stream ℬ, we use the previous *k* base call symbols as the context. Thus, for any given *k*-tuple *s* = *s*_1_*s*_2_ … *s*_*k*_, all symbols that follow an occurrence of *s* in ℬ are collected into the same context (we use a default value of *k* = 7). For the strand direction indicator stream, 𝒟, we directly encode each binary value *d*_*t*_ ∈ {0, 1} as a single bit, as experiments show that the strand direction indicators are close to uniformly distributed. The remaining streams are comprised of sequences of non-negative integers of various representation sizes. We use the same method to encode each such stream. To encode an *η*-bit integer stream, where *η* is a multiple of 8, we split the bit representation of each integer of the stream into 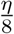 bytes. We encode each byte separately, starting from the least significant to the most significant byte, using a specific context for each of the 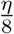 byte positions. In other words, all the least significant bytes of the integers in the stream are encoded in the same context, all the second least significant bytes are encoded in the same separate context, and so forth. Since, as mentioned, these non-negative integer sequences tend to concentrate towards the lower end of their range, the higher order bytes end up being highly compressible, and there is little penalty in over-estimating the selected integer size *η* for each stream.

#### 2.2.2 The decoding algorithm

In Algorithm 6 we present the general decompression scheme that decodes the sequence of base call strings *Q* = *q*_1_,…, *q*_|*Q*|_ given the set of reference base call strings ℛ and the compressed file, accessed through an input bitstream, *F*. Notice that, as the encoding and decoding algorithms work in lockstep, at any stage of the decoding process, the decoder knows exactly what stream to read from and how much information it needs from it.

Algorithm 6 reverses the steps of Algorithm 2. It starts by initializing *Q* as an empty sequence in Step 1 and all the streams in 𝕊 as empty streams in Step 2. The contents of both *Q* and 𝕊 are constructed in subsequent steps. First, the number of base call strings to be decoded and added to *Q, n*_*Q*_, is obtained from *F* in Step 3. Next, each stream in 𝕊 is decoded individually. This is done by retrieving the length, in bytes, of the encoding of the stream in Step 5, followed by the encoding itself in Step 6, which is decoded in Step 7. Once the streams in 𝕊 are decoded in Step 10, the algorithm reconstructs each of the read base call strings of *Q* from its stream representation by running Algorithm 7 with 𝕊 and ℛ as inputs. Next, we describe Step 10 in more detail.

##### Algorithm 6: Decode the sequence of read base call strings in a block of the FASTQ file.

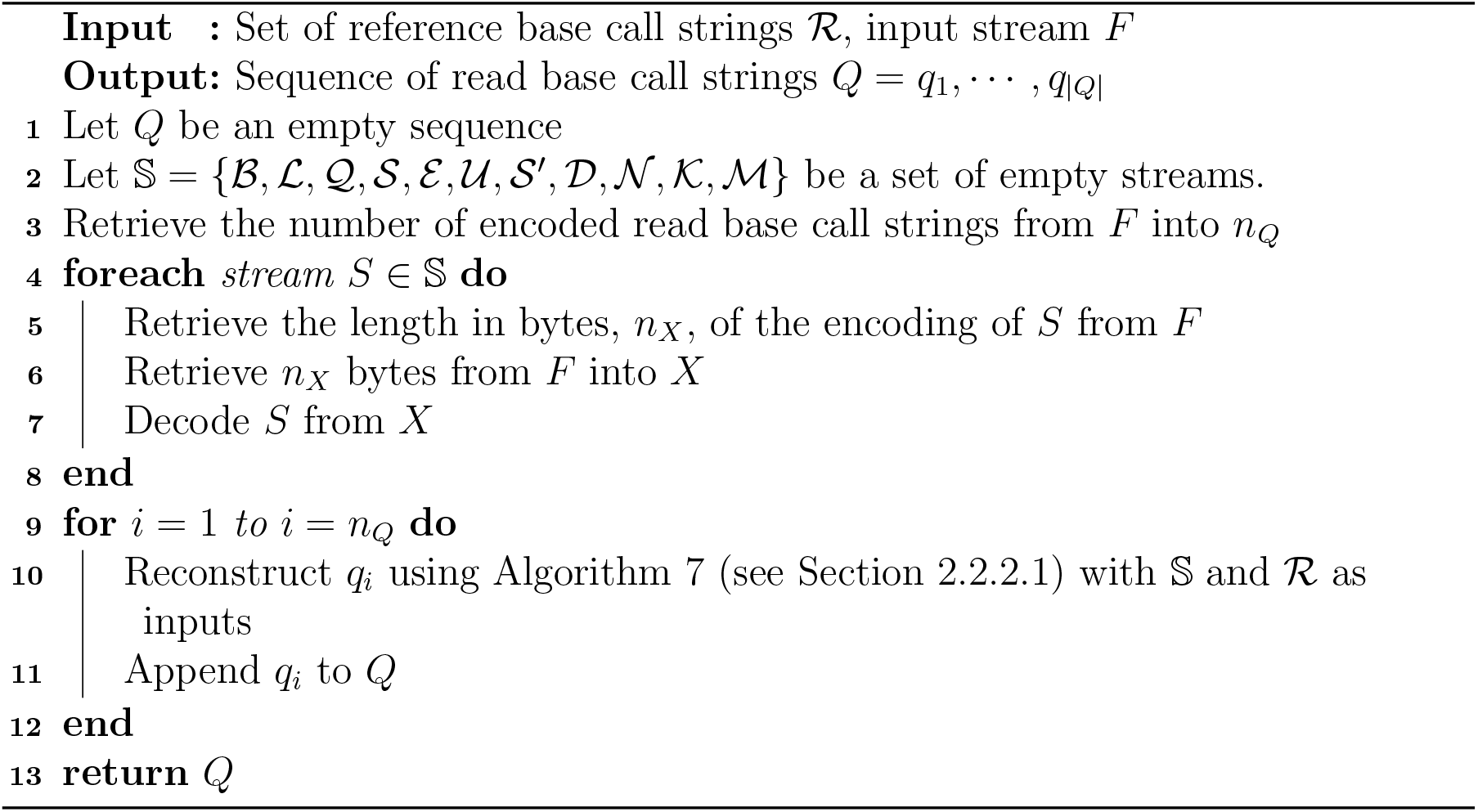

##### 2.2.2.1 Reconstructing read base call strings (Step 10 of Algorithm 6)

Algorithm 7 reconstructs a base call string *q* from its stream representation in 𝕊 and the set of reference strings ℛ.

Algorithm 7 reverses the steps of Algorithm 3. In the algorithm, *q* is initialized to an empty string in Step 1, and it is reconstructed progressively, appending symbols up to length *n*_*q*_ (obtained in Step 2). The variable *i*, initialized in Step 1, maintains the position of the next symbol to be added to *q*. In Step 3, the algorithm retrieves the number of atomic alignments *L*_*A*_ that encode substrings of *q*, and loops over each atomic alignment in Step 4. Each atomic alignment 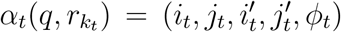 is retrieved in a two stage fashion. Firstly, *k*_*t*_ and *i*_*t*_ are obtained in Steps 5 and 6, respectively, of Algorithm 7. The index *i*_*t*_ is used to determine, in Steps 7–9, the length of an eventual unaligned string that may precede the atomic alignment. The remaining components of the atomic alignment, i.e., 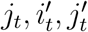, and *ϕ*_*t*_, are retrieved in a second stage by calling Algorithm 8 in step 10 with *i*_*t*_ and 𝕊 as inputs.

Algorithm 8 reverses the steps of Algorithm 4. Notice that, during the reconstruction process, we do not have direct access to the number of transformation steps, *K*, of the encoding transformation *ϕ*. However, the length of a string resulting from applying *ϕ* to a reference string equals the sum of the lengths of the insertion and match operations in *ϕ*. Therefore, for the length of the aligned substring, *z*, retrieved in Step 1, we have

#### Algorithm 7: Reconstruct base call string *q*.

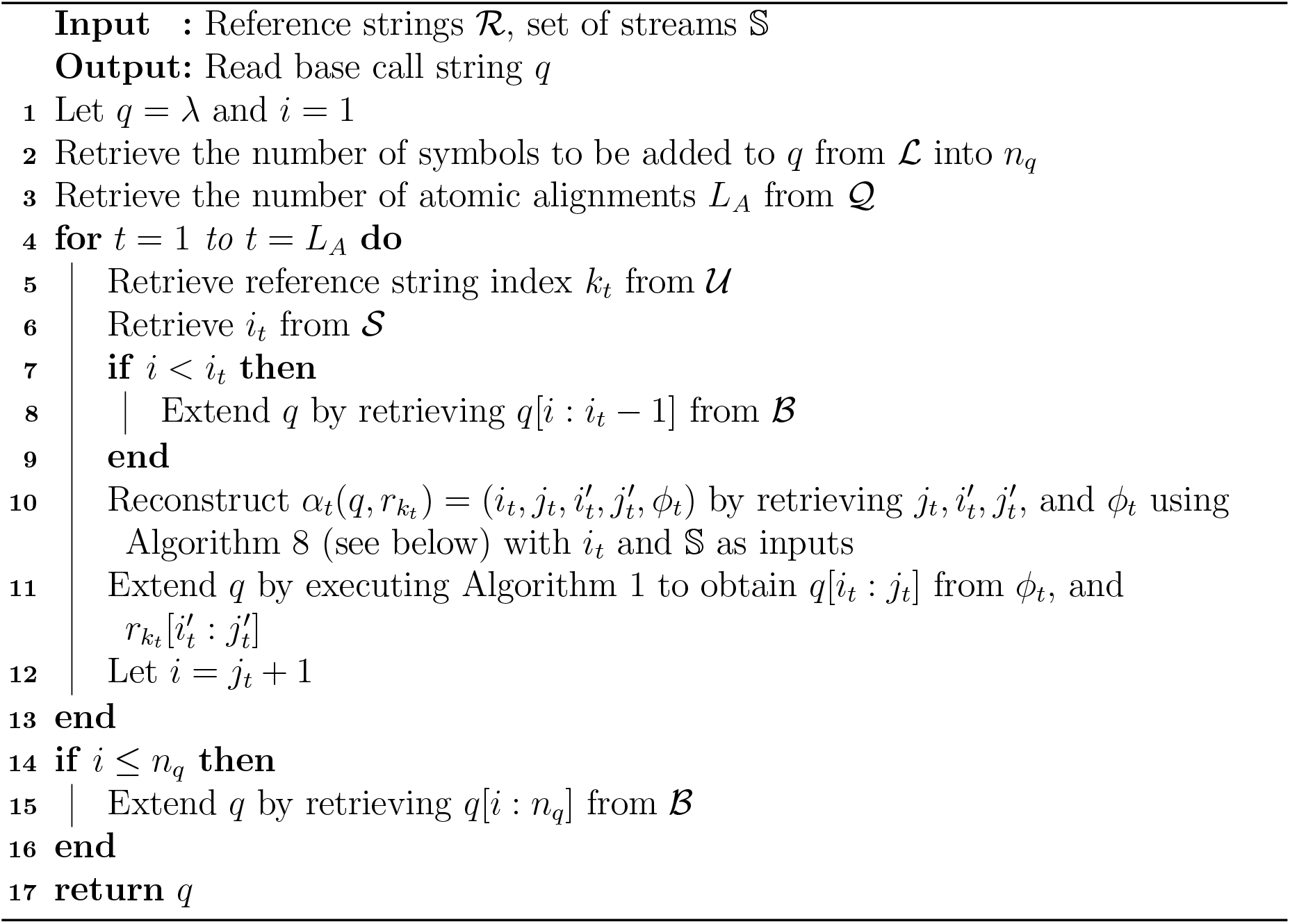

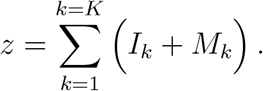

Consequently, the loop in Step 5 retrieves triplet operations, accumulating the lengths of the insertion and match operations in *z*_1_, until *z*_1_ reaches *z*.

###### Algorithm 8: Retrieve an atomic alignment from its stream representation in 𝕊.

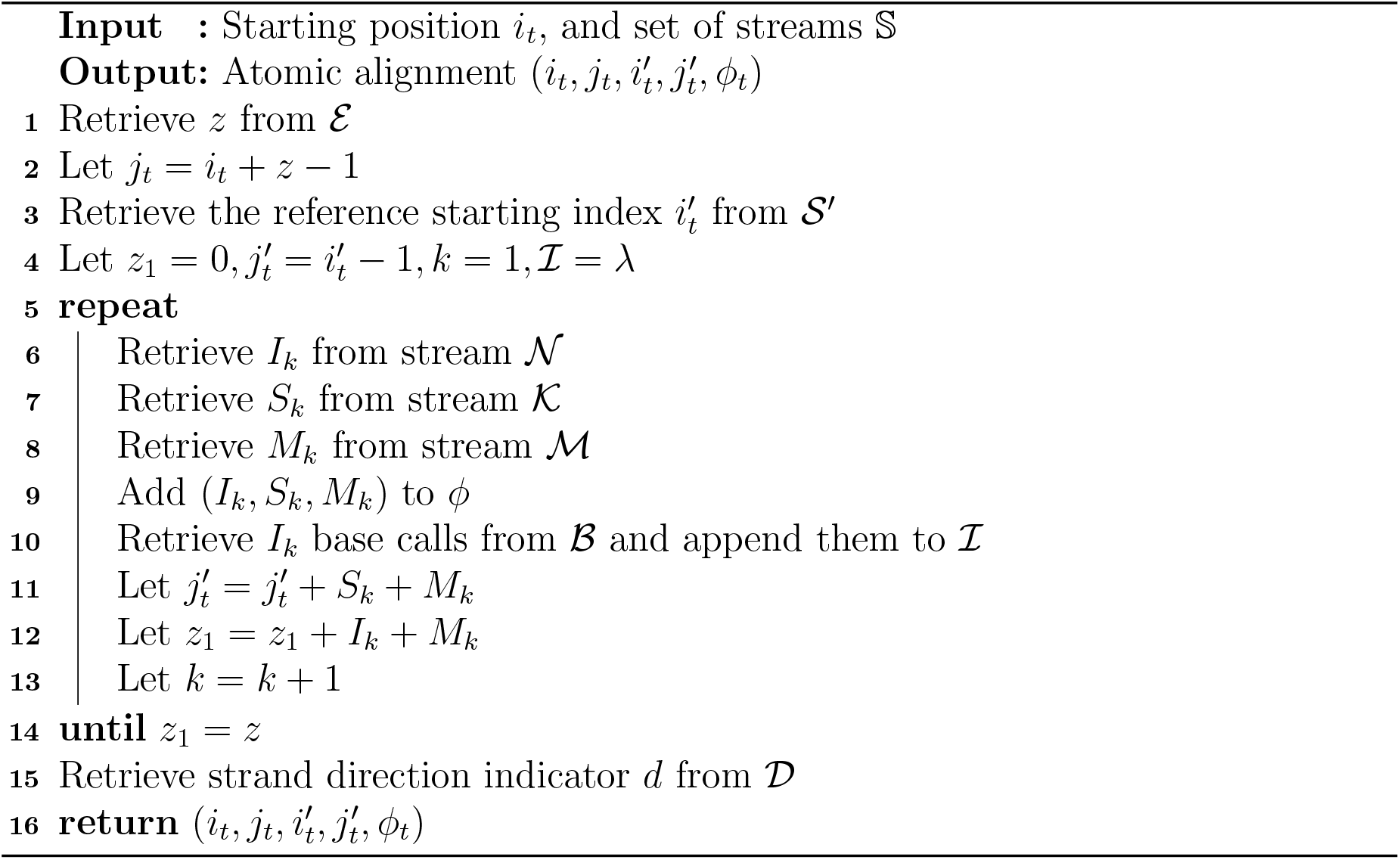

### 2.3 RENANO_2_ : a reference-dependent compression scheme, with a reference-independent decompression scheme

In this section we present RENANO_2_, a variation of RENANO_1_ that makes the decompression process independent of the set of reference base call strings ℛ. The main idea is to create a new artificial reference string, *r*′, composed of carefully selected parts of the set of reference base call strings ℛ and encode it in the compressed file. Briefly, starting from a preliminary alignment of the base call strings against ℛ, we construct *r*′ by retaining only parts of ℛ that are used by the atomic alignments, and hence that can help improving the compression. The remaining parts of ℛ are discarded since they are not needed by the decoder and therefore there is no need to include them in the compressed file. The atomic alignments associated to the read base call strings are then modified to align against *r*′ instead of the original strings in ℛ. At that point, we can compress the read base call strings of the FASTQ file by applying the same encoding scheme presented in Section 2.2.1, with ℛ = {*r*′}. Subsequently, we can decode the read base call strings by first decoding *r*′, followed by applying the decoding scheme of Section 2.2.2, again with ℛ = {*r*′}. Notice that, since ℛ consists of a single reference string, the index 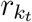 of each alignment is unnecessary and, thus, RENANO_2_ omits the encoding of the aligned reference string identity indexes stream 𝒰.

To create the new reference string *r*′, we start by analyzing the full alignments *A*(*q, ℛ*) obtained against the full reference set ℛ. We are interested in keeping the parts of the reference base call strings in ℛ that are deemed useful for the compression of the read base call strings *q*. More concretely, we first say that an atomic alignment, 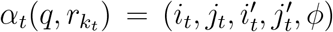, *uses* a position *h* of 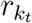 if 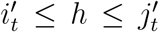. Naturally, the positions of a reference string *r*_*k*_ that are useful for compression are those that are actually used by atomic alignments, and we would like to discard positions that are not used by any atomic alignment. Moreover, taking into account that describing *r*′ incurs a coding cost, positions used by only one atomic alignment are not beneficial for compression either, as both the raw base call symbol for that position in the reference *r*′ and the alignment information need to be encoded. We empirically observe that good compression results are obtained by keeping the positions of the reference base call strings that are used by at least two atomic alignments, with no significant improvement for larger thresholds. Consequently, we say that *h* is a *surviving position* of *r*_*k*_ if it is used by at least two atomic alignments. We define the new reference *r*′ as the concatenation of the bases in surviving positions of the reference strings. Figure 3 shows an example of the proposed construction of *r*′.

**Figure 3:**
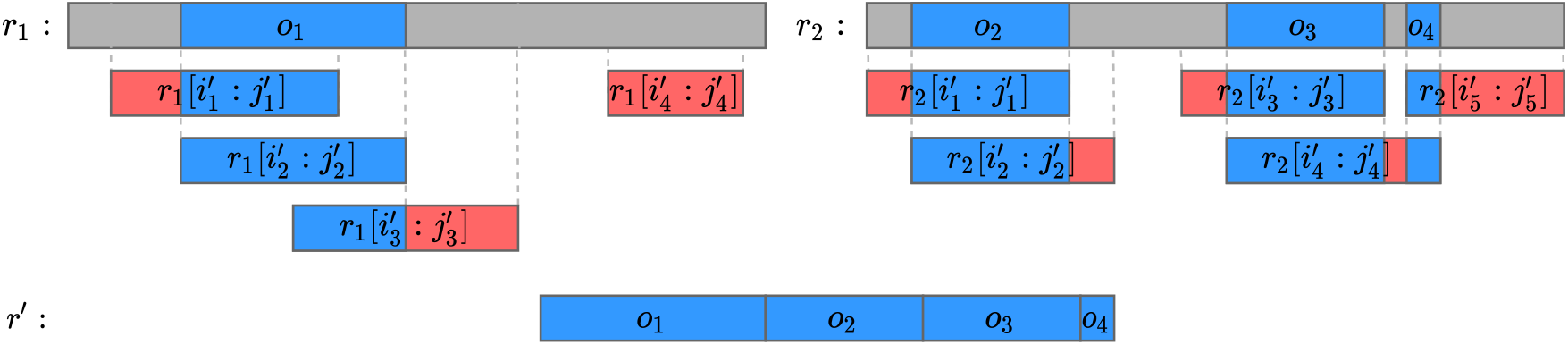
Example of the construction of an artificial reference string *r*′. The top row represents the original reference strings *r*_1_ and *r*_2_. The bottom row represents the constructed new reference *r*′. The three middle rows represent reference substrings, 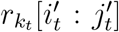, used by the atomic alignments. The string *r*′ is constructed by concatenating the bases in surviving positions (condensed in four segments, *o*_1_, *o*_2_, *o*_3_, and *o*_4_, marked in blue) of *r*_1_ and *r*_2_. Not-surviving positions (marked in gray) are discarded. In the reference substrings of the atomic alignments, the substrings that correspond to surviving positions are marked in blue, while substrings that correspond to not-surviving positions are marked in red.

Clearly, *r*′ can be constructed in a single pass through all atomic alignments, by keeping count of the number of uses of each position of *r*_*j*_, 1 ≤ *j* ≤. |ℛ | Moreover, as a byproduct of this construction, we can obtain a mapping *ψ* that maps each surviving position *h* of *r*_*j*_ to its corresponding position, *ψ*(*j, h*), in *r*′.

Once the new reference string *r*′ is built, RENANO_2_ generates a binary encoding, denoted by Enc(*r*′), using the same technique proposed for encoding the stream of base call strings ℬ in Section 2.2.1.2. The algorithm then stores Enc(*r*′) into the compressed file by outputting a fixed-size binary representation of its length | Enc(*r*′) |, followed by Enc(*r*′) itself. Clearly, this process can be reversed on the decoder side, giving the decoder access to *r*′.

To generate the artificial reference *r*′, we made use of the information in the atomic alignments, which were aligned against the original reference strings in ℛ. However, the decoder has access to *r*′, but not to ℛ. Therefore, before being described to the decoder, each atomic alignment, 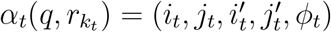, must be modified to align against *r*′. If the atomic alignment uses no surviving positions (see 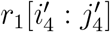 in Figure 3), it is fully discarded and removed from its respective full alignment. Otherwise, the atomic alignment is assigned a new reference substring of *r*′, 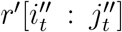, with 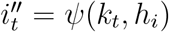 and 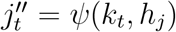, where *h*_*i*_ and *h*_*j*_ are smallest and largest surviving positions used by 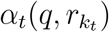, respectively. In addition, *ϕ*_*t*_ is modified so that 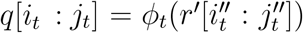, taking into account that some positions of the original reference substrings are no longer present in *r*′. Specifically, all triplet operations (*I*_*k*_, *S*_*k*_, *M*_*k*_) where a portion of the match or skip operations refers to not-surviving positions, are transformed: match operations are transformed into insertion operations, and the portions of skip operations that lie on not-surviving positions are removed. If during this process a triplet operation (*I*_*k*_, *S*_*k*_, *M*_*k*_) ends up with a value of *M*_*k*_ = 0, and it is not the last triplet of the encoding transformation, then the triplet is merged with the next triplet operation, (*I*_*k*+1_, *S*_*k*+1_, *M*_*k*+1_), resulting in a single triplet operation (*I*_*k*_ + *I*_*k*+1_, *S*_*k*_ + *S*_*k*+1_, *M*_*k*+1_).

At first sight, it may seem that the construction of *r*′ requires loading all the atomic alignments into memory at once. In practice, however, we can build *r*′ by loading, initially, only the information of the atomic alignments strictly required to determine the surviving positions, that is, the indexes of the aligned substrings *k*_*t*_, 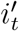 and 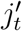. The remaining information of each atomic alignment is loaded only at the time of its encoding, where also the transformation of *ϕ*_*t*_ is performed.

### 2.4 Alignment Information

In this section, we describe the format in which the alignment information is provided to RENANO_1_ and RENANO_2_, and how we construct, for each base call string *q* in the FASTQ file, a full alignment *A*(*q, ℛ*) (as defined in Section 1, in our internal representation) against the set of reference sequences ℛ.

To obtain the alignment information of a specific FASTQ file against a reference genome, our software implementation of RENANO receives an input text file in PAF format, typically used for nanopore sequence alignment. Each line of a PAF file, which represents an alignment between a *query* base call string and a *target* base call string, consists of a list of TAB-delimited fields. For our application, each query base call string is a portion of a read base call string of a FASTQ file to be compressed, and each target base call string is a portion of a base call string of a reference genome, stored in a FASTA file. In Table 1 we describe the fields that are relevant to RENANO_1_ and RENANO_2_. Most fields are identified by position, with the only exception of the field labeled *cs*, which is identified through a tag at the beginning of the field.

**Table 1:** Fields of the PAF alignment format relevant to RENANO_1_ and RENANO_2_.

| Field | Type | Description |
| --- | --- | --- |
| 1 | string | Query string name |
| 3 | int | Query start coordinate |
| 4 | int | Query end coordinate |
| 5 | char | ‘+’ if query/target on the same strand; ‘-’ if opposite |
| 6 | string | Target string name |
| 8 | int | Target start coordinate on the original strand |
| 9 | int | Target end coordinate on the original strand |
| cs | string | Base-alignment string in *cs* format |

Our implementation reads and parses the input PAF file, line by line, to generate full alignments for each base call string *q* of the FASTQ file. The first field in each line of a PAF file is the sequence identifier of the read base call string in the FASTQ file. This read base call string, together with the starting and ending positions in fields 3 and 4 of the PAF file line, respectively, determine the query base call string of the alignment represented by this line. Each read base call string of a FASTQ file, *q*, may participate in zero or more alignments; each one is represented by a separate line in a PAF file, all of which bear the same sequence identifier of *q* in the first field. If a read base call string *q* has no associated alignments, then *A*(*q*, ℛ) is set to an empty list, and *L*_*A*_ = 0. Otherwise, for each line in the PAF file where *q* is the query string, we build an atomic alignment 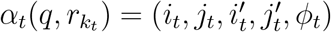. Recall from Section 2 that RENANO_1_ assumes that the atomic alignments of a full alignment *A*(*q*, ℛ) are non-overlapping, that is, that the aligned substrings, *q*[*i*_*t*_ : *j*_*t*_], of the atomic alignments in *Ā*(*q, ℛ*) do not overlap. However, this condition is not necessarily satisfied by the alignments extracted from the PAF file. Consequently, we first create an auxiliary full alignment *Ā*(*q*, ℛ) by parsing every atomic alignment associated to *q* in the PAF file, and we later use *Ā*(*q*, ℛ) to generate a non-overlapping full alignment *A*(*q*, ℛ).

To generate an atomic alignment 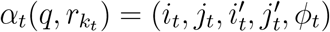 from a line of the PAF file, we use the values in fields 3 and 4 to determine the aligned *q* substring, *q*[*i*_*t*_ : *j*_*t*_], where *i*_*t*_ is the value in field 3 and *j*_*t*_ is the value in field 4. We also use the values in fields 6, 8, and 9, to determine the reference substring of the atomic alignment, 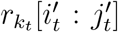, where the index of the reference substring *k*_*t*_ is determined by the name of the target string in field 6, and fields 8 and 9 determine 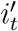 and 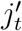, respectively. Finally, to determine the encoding transformation *ϕ*_*t*_, we use field 5 and the *cs* tag. Field 5 indicates if the alignment is between the original query string and the target string, in which case the value is ‘+’, or if it is between the reverse complement of the query string and the target string, in which case the value is ‘-’. We use this field to determine the strand direction indicator *d* of *ϕ*_*t*_, such that *d* = 0 if field 5 is ‘+’, and *d* = 1 if field 5 is ‘-’. Finally, the *cs* field has a base-alignment string, called *cs*-string, which encodes the difference between the query string and the target string, as a sequence of string operations, which include: *matches, insertions, substitutions* (single nucleotide polymorphisms), and *deletions*. For more information on *cs*-strings we refer the reader to the Supplementary Data of [20]. We use Algorithm 9 to process the *cs*-string, and obtain the triplet sequence {(*I*_*k*_, *S*_*k*_, *M*_*k*_)}_1≤*k*≤*K*_ and the insertions base call string, ℐ, of an encoding transformation *ϕ*_*t*_, such that 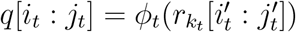.

The value of input *d* determines if the *cs*-string alignment utilizes the original query string, or its reverse complement. Thus, in Step 1, the algorithm starts by defining the auxiliary string *q*′ = *π*(*q*[*i*_*t*_ : *j*_*t*_], *d*), which is used to build the insertions base call string. In Step 2, the algorithm initializes the insertions base call string ℐ as the empty string, and the triplet sequence *T* as an empty sequence. The triplet operations and the insertions base call string are progressively constructed by scanning the *cs*-string and *q*′ from start to end as the algorithm iterates over the *cs*-string operations, appending triplets to *T* and base call strings to ℐ as necessary. The variable *i*, initialized to *i* = 1 in Step 3, maintains the starting position of the portion of *q*′ that remains to be processed. The auxiliary variable *I* represents the insertion length value of the next triplet to be appended, while *S* represents the skip length value of the next triplet to be appended. Both variables are initialized to 0 in Step 3.

In Step 4, the algorithm loops over each operation, *X*, in the *cs*-string, and takes a different action depending on the type of operation being processed. If the operation is an insertion, the algorithm adds the length of the operation, *L*, to the current insertion length, *I* (Step 7), and appends the corresponding inserted base calls, *q*′[*i* : *i* + *L* −1], to string ℐ (Step 8). The algorithm proceeds to adjust the value of index *i* accordingly (Step 9). If the operation is a deletion, the algorithm adds the deletion length *L* to the current skip length *S*, in Step 11. We interpret a substitution operation as a combination of an insertion and a deletion both of length 1. Therefore, for this kind of operations, the algorithm adds 1 to the value of *I* (Step 13), appends the base call in position *i* of *q, q*′[*i* : *i*], to ℐ (Step 14), and adds 1 to the current skip value *S* (Step 15). The index *i* is adjusted accordingly in Step 15. Finally, if the operation is a match, the algorithm checks, in Step 17, if the match length *L* is greater than a *minimum match length* threshold parameter, *m*_*L*_. If it is, the triplet (*I, S, L*) is appended to *T*, and the variables *I* and *S* are reset to 0, in steps 18 and 19, respectively. Otherwise, the match operation is interpreted as an insertion and a deletion, both of length *L*. Hence, the algorithm adds *L* to both *I* and *S* (Step 21), and appends string *q*′[*i* : *i* + *L* −1] to ℐ (Step 22). Index *i* is adjusted accordingly in Step 24. As a last step, if there is a remaining triplet operation that has yet to be appended to *T*, that is, if *I* or *S* are greater than zero, the corresponding triplet is appended to *T* (Step 28).

#### Algorithm 9: Obtain the sequence of triplet operations and the insertions base call string, from a *cs*-string, the aligned substring of *q*, and the strand direction indicator *d*.

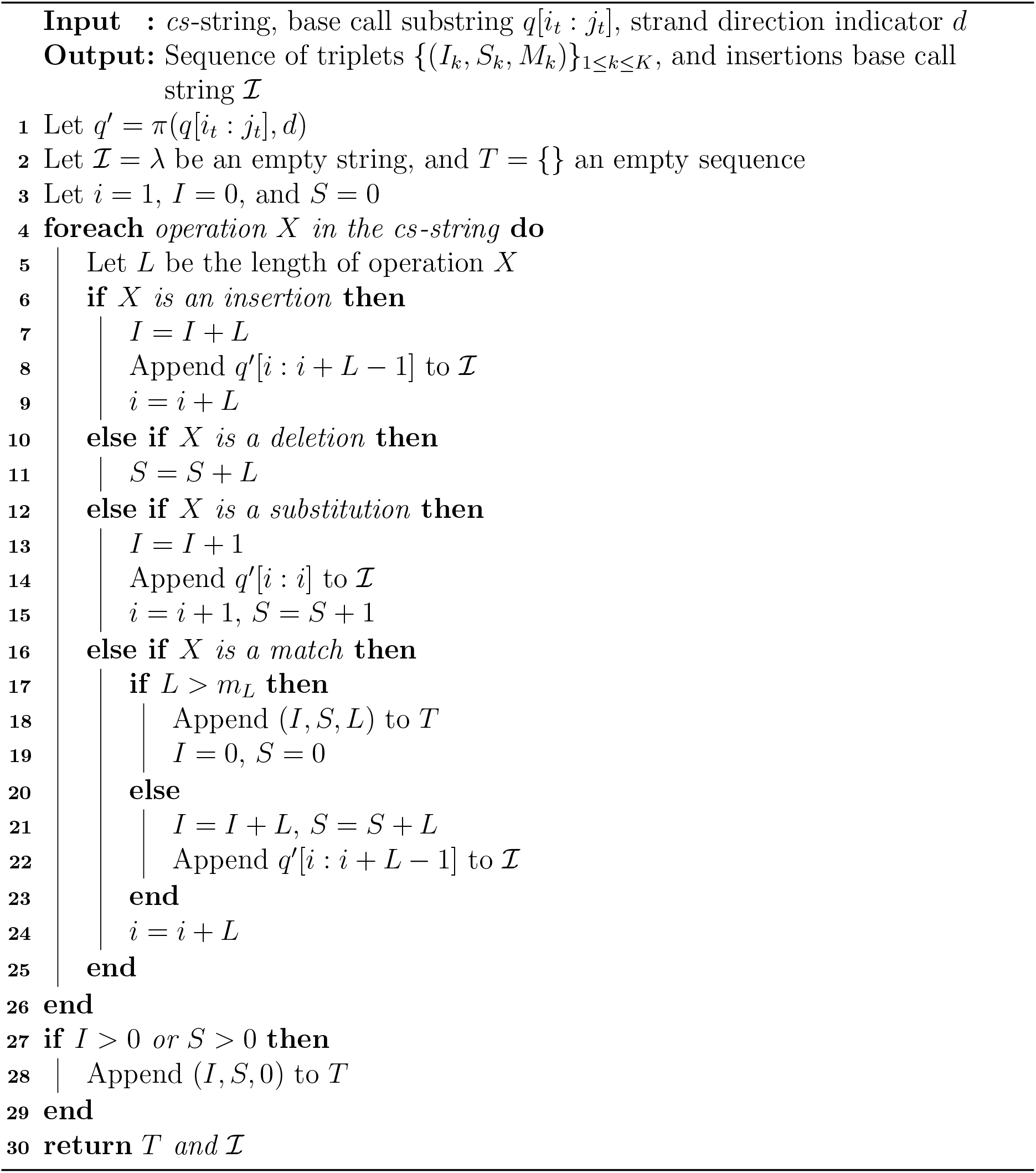

The parameter *m*_*L*_ regulates the trade off between the cost of encoding a potentially long sequence of atomized small matches, against the cost of encoding fewer insertion operations, which however incur an additional cost of encoding extra symbols in the insertions base call string. The effect of this parameter on compression performance is studied in Section 3.3, where we choose a default value for it.

Once the alignments in the PAF file associated to a base call string *q* are parsed and transformed into an auxiliary full alignment sequence 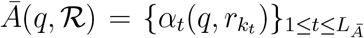, we generate a non-overlapping full alignment, *A*(*q*, ℛ), by executing Algorithm 10 with *Ā*(*q*, ℛ) as input.

The algorithm starts by sorting the sequence of atomic alignments, 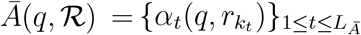, in decreasing order of *j*_*t*_ (Step 1). In Step 2, the sequence *A*(*q*, ℛ) is initialized with the first atomic alignment of the sorted sequence, 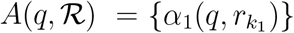. The algorithm progressively appends atomic alignments to *A*(*q*, ℛ), such that *A*(*q*, ℛ) always holds a non-overlapping sequence of atomic alignments. Notice that *A*(*q*, ℛ) is initialized as a non-overlapping sequence, as it contains a single atomic alignment. An example of a sequence of overlapping atomic alignments, sorted in decreasing order of *j*_*t*_, is presented in the top (pink) box in Figure 4.

**Figure 4:**
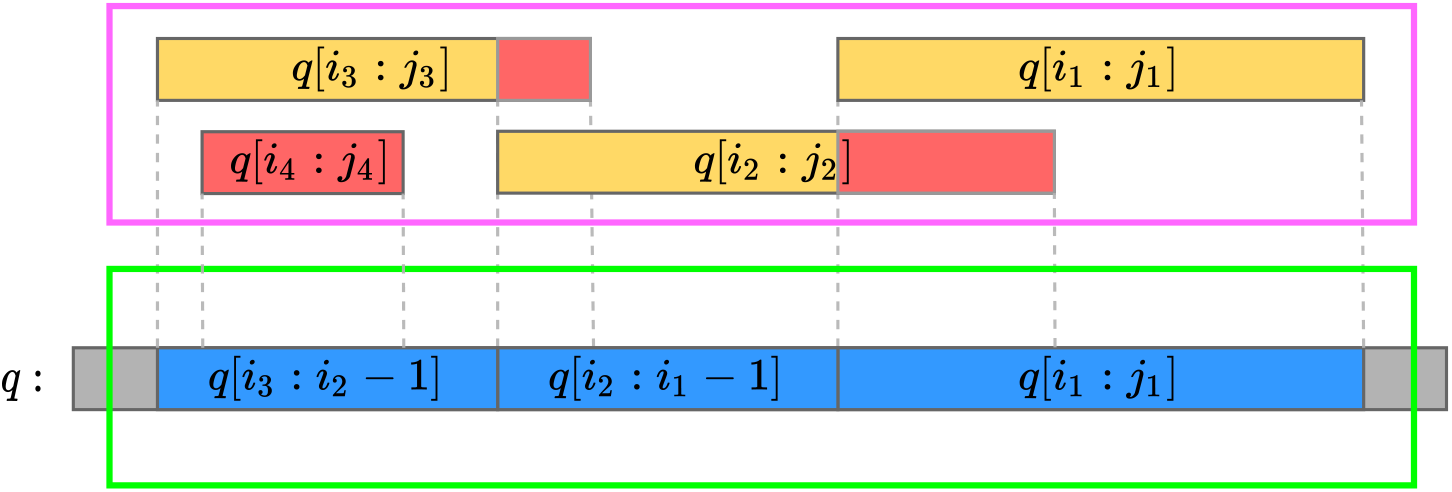
Example of a full alignment, 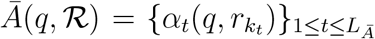, obtained from a PAF file with overlapping between aligned *q* substrings, *q*[*i*_*t*_ : *j*_*t*_]. The segments in the top box (pink) represent the original overlapping *q* substrings, while the segments in the bottom box (green) represent the result of running Algorithm 10 with *Ā*(*q*, ℛ) as input.

The variable *î*, initialized to *î* = *i*_1_ in Step 3 of the algorithm, maintains the starting position of the last atomic alignment appended to *A*(*q*, ℛ). In Step 4, the algorithm loops over the remaining atomic alignments in *Ā*(*q*, ℛ), and checks, in Step 5, if the ending position of the current atomic alignment, *j*_*t*_, is smaller than *î*. If so, the current alignment does not overlap with the alignments in *A*(*q*, ℛ) (as *i*_*t*_ < *j*_*t*_), and it is directly appended to *A*(*q*, ℛ) in Step 6, while *î* is updated to *î* = *i*_*t*_, in Step 7. Otherwise, if *j*_*t*_ is greater or equal than *î*, there is overlapping between *q*[*i*_*t*_ : *j*_*t*_] and *A*(*q*, ℛ). In this case, the algorithm checks, in Step 8, if *i*_*t*_ is smaller than *î*. If so, the prefix *q*[*i*_*t*_ : *î* − 1] of *q*[*i*_*t*_ : *j*_*t*_] does not overlap with *A*(*q*, ℛ) and, thus, an atomic alignment for this portion of *q* is added to *A*(*q*, ℛ) (see the yellow portions of the examples *q*[*i*_2_ : *j*_2_] and *q*[*i*_3_ : *j*_3_] in Figure 4). First, the variable *ĵ*_*t*_ is initialized to *ĵ*_*t*_ = *î* − 1 in Step 9, which is the ending of the non-overlapping portion of *q*[*i*_*t*_ : *j*_*t*_]. The algorithm then proceeds to construct 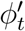, such that 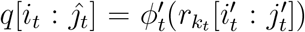, in Step 10, by modifying *ϕ*_*t*_. Specifically, the parts of the triplet sequence and the insertions base call string of the encoding transformation *ϕ*_*t*_ that operate from position *î* to *j*_*t*_ are discarded. In Step 11, the modified version of the atomic alignment, 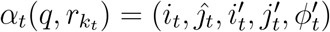, is appended to *A*(*q*, ℛ), and *î* is updated to *î* = *i*_*t*_ in Step 12. Notice that, if *i*_*t*_ is greater or equal than *î*, i.e., if the condition in Step 8 is not satisfied, then *q*[*i*_*t*_ : *j*_*t*_] fully overlaps with *A*(*q*, ℛ) in which case it is discarded (see example *q*[*i*_4_ : *j*_4_] in Figure 4). Finally, *A*(*q*, ℛ) is returned in Step 15. In Figure 4, the bottom (green) box shows the result of executing Algorithm 10 with the auxiliary full alignment in the top (pink) box as input.

#### Algorithm 10: Generate a non-overlapping full alignment *A*(*q*, ℛ).

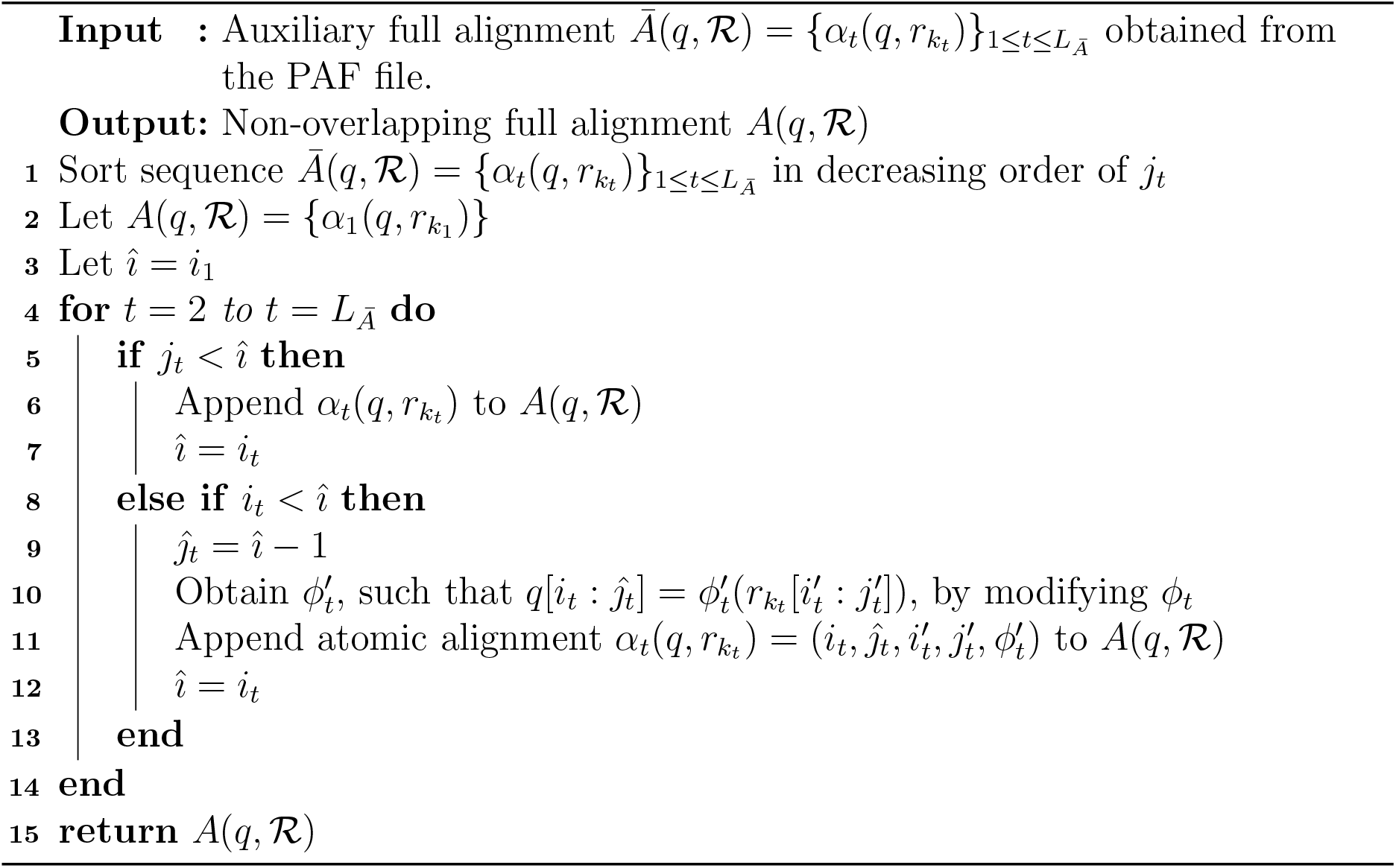

## 3 Experimental results

In this section we report on experiments performed on a collection of datasets of nanopore FASTQ files. We describe the datasets in Section 3.1. In Section 3.2 we explain how we generate the input PAF files that are necessary to run the proposed compression algorithms. In Section 3.3 we discuss the default values used for various algorithm parameters of RENANO_1_ and RENANO_2_. Finally, in Section 3.4 we evaluate the performance of RENANO_1_ and RENANO_2_ by comparing them against other compression tools.

To measure the performance of a compressor on a dataset, we compress each file of the dataset separately and calculate the *compression ratio, CR*, defined as *CR* = *C/T*, where *T* is the total size in bytes of the files (or file parts) under consideration, and *C* is the total size in bytes of the corresponding compressed files (or parts). Notice that smaller values of *CR* correspond to better compression performance. We will use the measure *CR* in two settings: in the first one, *C* and *T* refer to the base call strings part of the FASTQ file only, while in the second one *C* and *T* refer to the full FASTQ file. Thus, the first setting focuses only on the parts that are different in ENANO and RENANO, while the second setting reflects the impact on the overall compression of the FASTQ files. To compare compression ratios, we define the *percentage relative* 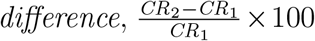, between the compression ratios *CR*_1_ and *CR*_2_, with respect to *CR*_1_. Clearly, negative values of this measure indicate that *CR*_2_ is better than *CR*_1_, while positive values indicate that *CR*_2_ is worse. Finally, to compare overall performance between compressors we report *simple* and *weighted* averages of the results over the test datasets, the latter computed by weighting each result by the size of its corresponding dataset. The weighted average is highly influenced by larger datasets and, in particular, by the specific choice of datasets available for evaluation. This can be misleading when assessing the capability of the tools to process different types of datasets. Hence, in the rest of this section we report both averages, but we generally refer to the simple average when discussing and comparing the evaluated tools, unless otherwise specified.

All experiments were conducted on a server with 80 64 bit x86 Intel Xeon CPUs, 503.5GB of RAM memory, and CentOS Linux release 7.7.1908.

### 3.1 Datasets

We evaluate the proposed algorithms on a collection of publicly available datasets, described in Table 2. The collection includes a metagenomic dataset (last entry in the table).

**Table 2:** Nanopore sequencing datasets used for evaluation.

| Name | Num. Files | Total size (GB) | Description |
| --- | --- | --- | --- |
| *hss* | 1 | 268 | Human GM12878 Utah/Ceph cell line [11] |
| *bra* | 18 | 46 | Brassica napus L. [22] |
| *sor* | 4 | 133 | Sorghum bicolor Tx430 [4] |
| *fly* | 1 | 17 | Drosophila ananassae [23] |
| *yst* | 5 | 6 | Saccharomyces cerevisiae S288C [10] |
| *mic* | 1 | 12 | Microbial community (metagenomic) [18] |

The selected datasets cover a variety of dissimilar organisms including human, plant, animal, fungi, and bacteria. In the case of non-metagenomic datasets, for each dataset we obtained a reference genome file from the NCBI database [24], from which we extract the reference base call strings used in our algorithms. The metagenomic bacterial dataset, *mic*, demonstrates a scenario in which we do not know in advance the species that are present in the sequenced samples. For this case, we propose a pipeline of operations for constructing a reference sequence. It consists of performing a taxonomic classification of the dataset reads, and then concatenating the reference genomes of the most prevalent organisms in the dataset. Several tools are available for the taxonomic classification step, such as FALCON [26], Kraken2 [30], and Centrifuge [13]. To facilitate the execution of this pipeline, we provide a set of scripts that automatize the reference sequence construction process at https://github.com/guilledufort/RENANO. These scripts identify the most prevalent species in a FASTQ file by running and analyzing the output of Kraken2, download the corresponding reference genomes directly from the NCBI database, and concatenate these genomes into a single FASTA file, which serves as the reference sequence for compression.

In Table 3 we present the total size of the read base call strings in each dataset, the identification string of the reference genome file associated to the dataset (except for the metagenomic dataset), the total size of the reference base call strings of the genome, and the coverage of the dataset (total and average per file).

**Table 3:**
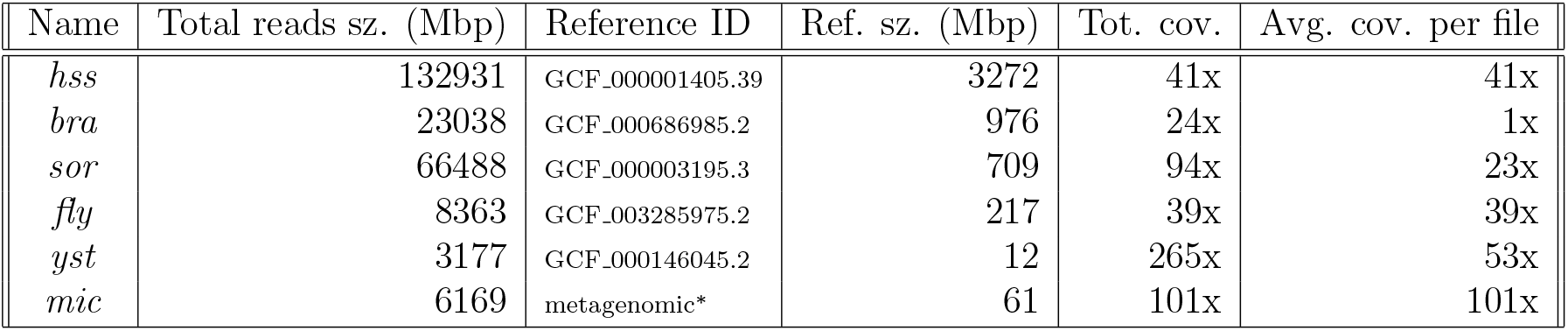
Read base call string information of the different datasets and their associated reference genomes. The total sizes of the read base call strings and the reference strings are presented in mega base-pairs (Mbp). The total coverage is calculated by dividing the total number of base call symbols in the dataset reads by the number of base call symbols in the strings of the reference genome. The average coverage per file was calculated by dividing the total coverage of the dataset by the corresponding number of files. ^∗^The reference sequence used for the microbial metagenomic dataset was constructed by first performing a taxonomic classification of the dataset reads, followed by concatenating the references of the most prevalent organisms.

The datasets cover different possible compression scenarios, such as having 268 GB of human data in a single file (hss), with 41x coverage, or having 46 GB of plant data distributed in 18 files (bra), with 1x average coverage per file, or having 12 GB of microbial metagenomic data in a single file, with 101x coverage. Further instructions on how to download the datasets, and the reference genomes, including instructions on how to construct a reference sequence for a metagenomic or contaminated dataset are available at https://github.com/guilledufort/RENANO.

### 3.2 PAF files generation

For our experiments, we generate PAF files with Minimap2 [20], a state of the art sequence alignment tool for long reads. We run Minimap2 with the FASTQ file we want to compress and the proper reference genome file as inputs. In turn, the tool outputs the result of aligning each base call string of the FASTQ file against the reference genome, in PAF format. Specifically, we execute Minimap2 with the following configuration options:

- *-x map-ont* : an option that sets the configuration of the tool to be optimized for reads generated with nanopore technologies. This option is recommended.
- *--cs*: an option that makes the tool perform base-alignment between the aligned sections of the read base call strings and the reference strings. The base-alignment is expressed as a *cs-string*. This option is necessary, as the base-alignment information is needed for our algorithms.
- *--secondary=no*: an option that makes the tool output only primary alignments. This prevents the tool from outputting multiple alignments for the same sections of the read base call strings. This option is recommended.

### 3.3 Algorithm parameters

In this section we specify the values of the parameters used in our implementations. First, we address the sizes of integers in the streams defined in Section 2.2.1. The following values were used in our experiments, but can eventually be modified as they are easily re-configurable: *η*_ℒ_ = 24, *η*_𝒬_ = 8, *η*_𝒮_ = 24, *η*_ℰ_ = 24, *η*_𝒰_ = 16, *η*_𝒮 ′_ = 32, *η*_𝒩_ = 16, *η*_𝒦_ = 16, and *η*_ℳ_ = 16.

Furthermore, in Algorithm 9, we introduce the minimum match length threshold, *m*_*L*_. This parameter determines which match operations in the base-alignment *cs*-strings are interpreted as match operations when transformed into the triplets of encoding transformations described in Section 2.1, and which ones are disregarded and interpreted as an insertion. To determine the default value of *m*_*L*_, first we evaluate its impact on the compression performance of RENANO_1_, by running the algorithm with different values of *m*_*L*_, ranging from *m*_*L*_ = 1 to *m*_*L*_ = 6, on the datasets specified in Table 2. The compression results for the base call strings of each dataset are shown in Table 4.

**Table 4:** Reads base call strings compression ratio of RENANO_1_, with different values of minimum match length *m*_*L*_. Best results for each dataset are bold-faced.

| Dataset | $m_z = 1$ | $m_z = 2$ | $m_z = 3$ | $m_z = 4$ | $m_z = 5$ | $m_z = 6$ |
| --- | --- | --- | --- | --- | --- | --- |
| *hss* | 0.118 | **0.117** | 0.118 | 0.119 | 0.120 | 0.122 |
| *bra* | **0.141** | **0.141** | 0.142 | 0.143 | 0.145 | 0.147 |
| *sor* | 0.161 | **0.160** | 0.161 | 0.162 | 0.164 | 0.167 |
| *fly* | 0.118 | **0.117** | 0.118 | 0.119 | 0.121 | 0.123 |
| *yst* | 0.174 | **0.173** | **0.173** | 0.174 | 0.175 | 0.177 |
| *mic* | **0.158** | **0.159** | 0.160 | 0.161 | 0.162 | 0.164 |

The results show that the best compression performance for most of the tested datasets is obtained at *m*_*L*_ = 2, which we set as the default value in RENANO_1_ and RENANO_2_ (the minimum is rather shallow, though, so the precise choice of value is not critical).

### 3.4 Comparative experimental results

In this section we evaluate RENANO_1_ and RENANO_2_ by comparing their performance against other compression tools. In Section 3.4.1, we compare against the state of the art nanopore sequencing FASTQ data compressor ENANO [5], on the datasets specified in Table 2. In Section 3.4.2, we also compare against the FASTQ compressor Genozip [16], a recently developed compression tool that can be used for nanopore data and offers both reference-based and reference-free compression modes.

#### 3.4.1 Comparison against ENANO

To evaluate the performance of RENANO_1_ and RENANO_2_ against ENANO, we run each compressor on the datasets specified in Table 2. Each compressor is configured to run in its default configuration (see Section 3.3 for RENANO and [5] for ENANO). The read base call strings and total compression ratios obtained on each dataset are shown in Table 5.

**Table 5:** Read base call strings and total compression ratios (*CR*) for ENANO, RENANO_1_, and RENANO_2_, on all the datasets. The percentage relative difference of RENANO_1_, and RENANO_2_ with respect to ENANO are shown in parenthesis. The table also shows the simple (S.) and weighted (W.) *CR* averages. Best results for each dataset are bold-faced.

| Dataset | Read base call strings |  |  | Total |  |  |
| --- | --- | --- | --- | --- | --- | --- |
|  | ENANO | RENANO<sub>1</sub> | RENANO<sub>2</sub> | ENANO | RENANO<sub>1</sub> | RENANO<sub>2</sub> |
| *hss* | 0.236 | **0.117 (-50.3)** | 0.123 (-47.8) | 0.452 | **0.393 (-13.0)** | 0.396 (-12.4) |
| *bra* | 0.238 | **0.141 (-40.7)** | 0.202 (-15.2) | 0.355 | **0.307 (-13.6)** | 0.337 (-5.1) |
| *sor* | 0.245 | **0.160 (-34.6)** | 0.171 (-30.3) | 0.374 | **0.332 (-11.3)** | 0.337 (-9.9) |
| *fly* | 0.242 | **0.117 (-51.5)** | 0.123 (-49.0) | 0.354 | **0.292 (-17.3)** | 0.295 (-16.5) |
| *yst* | 0.236 | **0.173 (-26.9)** | 0.178 (-24.7) | 0.297 | **0.266 (-10.4)** | 0.269 (-9.6) |
| *mic* | 0.244 | **0.159 (-35.0)** | 0.162 (-33.8) | 0.407 | **0.364 (-10.5)** | 0.365 (-10.2) |
| *S. average* | 0.240 | **0.145 (-39.8)** | 0.160 (-33.5) | 0.373 | **0.326 (-12.7)** | 0.333 (-10.6) |
| *W. average* | 0.239 | **0.133 (-44.4)** | 0.146 (-39.2) | 0.415 | **0.362 (-12.6)** | 0.368 (-11.1) |

The results show that both RENANO_1_ and RENANO_2_ outperform ENANO for all the datasets. In particular, the best result for each dataset is achieved by RENANO_1_. For read base call strings compression, RENANO_1_ shows improvements relative to ENANO ranging from 26.9% (in *yst*) to 51.5% (in *fly*), with an average improvement of 40.8% over all the datasets. As for total compression, the improvements range from 10.4% (in *yst*) to 17.3% (in *fly*), with an average improvement of 13.1%. RENANO_2_ also consistently improves over ENANO. For read base call strings, the improvements range from 15.2% (in *bra*) to 49.0% (in *fly*), while in terms of total size, the improvements range from 5.1% (in *bra*) to 16.5% (in *fly*).

Compared to RENANO_1_, RENANO_2_ achieves similar compression results in the datasets with high coverage per file (*hss* 41x, *sor* 23x, *fly* 39x, and *yst* 53x), with a relative percentage deterioration with respect to RENANO_1_ ranging from 2.9% (in *yst*) to 6.9% (in *sor*). In the case of dataset *bra*, where the average coverage per file is 1x, the relative percentage deterioration between RENANO_2_ and RENANO_1_ reaches 43.3%. This is due to RENANO_2_ directly benefiting from having multiple atomic alignments that use the same sections of the reference strings, which is less likely to happen in files with low coverage. However, even for the dataset *bra*, which has an average coverage per file as low as 1x, RENANO_2_ improves the compression of the read base call strings by 15.2% relative to ENANO, which leads to a total compression improvement of 5.1%. We also notice that for the metagenomic dataset, *mic*, for which we constructed a reference sequence following the pipeline described in Section 3.1, both RENANO_1_ and RENANO_2_ improve the base call strings compression performance of ENANO by 35.0% and 33.8%, respectively.

In Tables 6 and 7 we show the total compression and decompression times (in *h:mm:ss* format) and speeds (in MB/s), respectively, for each algorithm on each dataset. All the algorithms were run in multi-threading environments, using eight threads in each run.

**Table 6:** Encoding and decoding times (in *h:mm:ss* format) for ENANO, RENANO_1_, and RENANO_2_, on all the datasets. Best results, for each dataset, both for encoding and decoding, are bold-faced.

| Dataset | Total compression time (h:mm:ss) |  |  | Total decompression time (h:mm:ss) |  |  |
| --- | --- | --- | --- | --- | --- | --- |
|  | ENANO | RENANO<sub>1</sub> | RENANO<sub>2</sub> | ENANO | RENANO<sub>1</sub> | RENANO<sub>2</sub> |
| *hss* | **0:37:09** | 0:49:56 | 0:51:24 | 0:52:55 | **0:49:40** | 0:51:19 |
| *bra* | **0:05:26** | 0:11:32 | 0:08:30 | **0:07:40** | 0:07:52 | 0:09:34 |
| *sor* | **0:13:09** | 0:14:51 | 0:17:34 | 0:19:45 | **0:19:15** | 0:20:33 |
| *fly* | **0:01:41** | 0:03:23 | 0:02:08 | 0:02:30 | **0:02:19** | 0:02:26 |
| *yst* | **0:00:49** | 0:01:33 | 0:01:00 | **0:01:05** | **0:01:05** | 0:01:06 |
| *mic* | **0:01:16** | 0:01:20 | 0:01:33 | 0:01:53 | **0:01:48** | 0:01:51 |

**Table 7:** Encoding and decoding speeds in MB/s for ENANO, RENANO_1_, and RENANO_2_, on all the datasets. The table also shows the simple (S.) and weighted (W.) averages of the results. Best results, for each dataset, both for encoding and decoding, are bold-faced.

| Dataset | Compression speed (MB/s) |  |  | Decompression speed (MB/s) |  |  |
| --- | --- | --- | --- | --- | --- | --- |
|  | ENANO | RENANO<sub>1</sub> | RENANO<sub>2</sub> | ENANO | RENANO<sub>1</sub> | RENANO<sub>2</sub> |
| *hss* | **120** | 90 | 87 | 84 | **90** | 87 |
| *bra* | **142** | 67 | 91 | **101** | 98 | 81 |
| *sor* | **169** | 150 | 127 | 113 | **116** | 108 |
| *fly* | **168** | 83 | 132 | 113 | **122** | 116 |
| *yst* | **131** | 70 | 107 | 99 | **100** | 97 |
| *mic* | **162** | 153 | 133 | 109 | **114** | 111 |
| *S. average* | **149** | 102 | 113 | 103 | **107** | 100 |
| *W. average* | **139** | 105 | 101 | 95 | **100** | 94 |

The results show that ENANO is the fastest compressor for all the datasets, while RENANO_1_ is on average the fastest decompressor. Specifically, during compression ENANO was 1.3x and 1.4x times faster on average than RENANO_1_ and RENANO_2_, respectively. During decompression, RENANO_1_ was 1.05x and 1.1x times faster on average than ENANO and RENANO_2_, respectively.

In table 8 we show the maximum memory required by each compressor during the encoding and decoding processes on all the files of each dataset.

**Table 8:** Maximum memory usage (in GB) registered during the encoding and decoding processes, for all the compressors, on all the files of each dataset. Lowest memory usage, for each dataset, both for encoding and decoding, are bold-faced.

| Dataset | Max. compression memory use (GB) |  |  | Max. decompression memory use (GB) |  |  |
| --- | --- | --- | --- | --- | --- | --- |
|  | ENANO | RENANO<sub>1</sub> | RENANO<sub>2</sub> | ENANO | RENANO<sub>1</sub> | RENANO<sub>2</sub> |
| *hss* | **0.220** | 3.723 | 10.399 | **0.231** | 3.441 | 3.790 |
| *bra* | **0.210** | 1.187 | 1.539 | **0.226** | 1.182 | 0.868 |
| *sor* | **0.214** | 0.943 | 2.998 | **0.227** | 0.923 | 0.933 |
| *fly* | **0.210** | 0.441 | 1.134 | **0.225** | 0.439 | 0.410 |
| *yst* | **0.205** | 0.240 | 0.345 | **0.223** | 0.238 | 0.238 |
| *mic* | **0.342** | 0.432 | 0.701 | **0.304** | 0.370 | 0.345 |
| *Maximum* | **0.342** | 3.723 | 10.399 | **0.304** | 3.441 | 3.790 |

The results show that ENANO is the most efficient in terms of memory for all considered datasets, both for compression and decompression. This is in part due to RENANO_1_ and RENANO_2_ having to load a reference genome file into memory during the encoding and decoding processes (in the case of RENANO_2_ the artificial reference is loaded during decoding). Also, in the case of RENANO_2_, to generate the artificial reference during encoding, the algorithm needs to load information of each atomic alignment obtained from the PAF file, which makes the memory usage grow with the number of alignments. However, even for the alignment of the *hss* file of 286 GB, the memory usage is of 10.4 GB, which is manageable by a personal computer with 16 GB of RAM.

#### 3.4.2 Comparison against Genozip

We compare RENANO_1_ and RENANO_2_ against the tool Genozip [16], which has two modes: a reference-free compression mode (which we refer a to as *Gen*), and a reference-based compression mode (which we refer to as *Gen-ref*). For this comparison we run the selected tools on the nanopore datasets specified in Table 2. We execute both modes of Genozip configured to maximize compression (i.e., in their default configurations). In the case of the reference-based compression mode, we use the reference genome files presented in Table 3. In Table 9 we present the read base call strings, and total, compression ratios obtained by running both modes of Genozip, RENANO_1_, and RENANO_2_, on the selected datasets.

**Table 9:** Read base call strings, and total, compression ratios for both modes of Genozip, RENANO_1_, and RENANO_2_, on all the datasets. The percentage relative difference of RENANO_1_ and RENANO_2_ with respect to *Gen* is presented in parenthesis. Best results for each dataset are bold-faced. The table also shows the simple (S.) and weighted (W.) averages of the results. The symbol ‘-’ indicates the tool failed to compress the dataset.

| File from | Read base call strings |  |  |  | Total |  |  |  |
| --- | --- | --- | --- | --- | --- | --- | --- | --- |
|  | Gen | Gen-ref. | RENANO<sub>1</sub> | RENANO<sub>2</sub> | Gen | Gen-ref. | RENANO<sub>1</sub> | RENANO<sub>2</sub> |
| *hss* | 0.233 | - | **0.117 (-49.5)** | 0.123 (-46.9) | 0.476 | - | **0.393 (-17.5)** | 0.396 (-16.9) |
| *bra* | 0.236 | 0.282 | **0.141 (-40.2)** | 0.202 (-14.5) | 0.370 | 0.400 | **0.307 (-17.2)** | 0.337 (-9.0) |
| *sor* | 0.244 | - | **0.161 (-34.0)** | 0.171 (-30.1) | 0.390 | - | **0.332 (-14.8)** | 0.337 (-13.6) |
| *fly* | 0.244 | 0.282 | **0.117 (-51.9)** | 0.123 (-49.4) | 0.385 | 0.400 | **0.292 (-24.0)** | 0.295 (-23.2) |
| *yst* | 0.236 | 0.286 | **0.173 (-27.0)** | 0.178 (-24.8) | 0.313 | 0.339 | **0.266 (-14.8)** | 0.269 (-14.0) |
| *mic* | 0.244 | 0.281 | **0.159 (-34.9)** | 0.162 (-33.7) | 0.417 | 0.435 | **0.364 (-12.6)** | 0.365 (-12.3) |
| *S. average* | 0.240 | 0.283 | **0.145 (-39.6)** | 0.160 (-33.2) | 0.392 | 0.394 | **0.326 (-16.8)** | 0.333 (-14.8) |
| *W. average* | 0.237 | 0.282 | **0.133 (-43.7)** | 0.146 (-38.6) | 0.435 | 0.400 | **0.362 (-16.8)** | 0.368 (-15.3) |

With respect to the results obtained by the two modes of Genozip, the reference-free mode achieved better results than the reference-based mode in all the datasets. Note also that the reference-based mode failed to compress the *hss* and the *sor* datasets. Therefore, to evaluate the performance of RENANO_1_ and RENANO_2_ we compare them against the reference-free mode of Genozip (Gen).

The results show that both RENANO_1_ and RENANO_2_ significantly outperform Genozip in each of the datasets. Specifically, regarding read base call strings compression, RENANO_1_ and RENANO_2_ improve on Genozip by 40.5% and 33.1% on average, respectively, and in total compression by 17.7% and 15.3% on average, respectively.

